# Metabolic potentials of *Liquorilactobacillus nagelii* AGA58 isolated from Shalgam based on genomic and functional analysis

**DOI:** 10.1101/2021.12.12.472281

**Authors:** Ahmet Yetiman, Fatih Ortakci

**Affiliations:** Erciyes University, Faculty of Engineering, Food Engineering Department, Kayseri, Turkey; Abdullah Gül University, Faculty of Life and Natural Sciences, Department of Bioengineering, Kayseri, Turkey

**Keywords:** *Liquorilactobacillus nagelii*, genome, motile, homofermentative, probiotic

## Abstract

The aim of present study was to perform functional and genomic characterization of a novel *Liquorilactobacillus nagelii* AGA58 isolated from Shalgam to understand its metabolic potentials. AGA58 is gram-positive, catalase-negative and appears as short-rods under light-microscope. The AGA58 chromosome composed of a single linear chromosome of 2,294,535 bp that is predicted to carry 2151 coding sequences, including 45 tRNA genes, 4 rRNA operons. Genome has a GC content of 36.9% includes 45 pseudogenes, 32 transposases and one intact-prophage. AGA58 is micro-anaerobic owing to shorter doubling time and faster growth rate achieved compared microaerofilic condition. It carries flagellar biosynthesis protein-encoding genes predicting motile behavior. AGA58 is an obligatory homofermentative where hexose sugars such as galactose, glucose, fructose, sucrose, mannose, N-acetyl glucosamine, maltose, trehalose are fermented to lactate thru glycolysis and no acid production from pentose sugars achieved due to lack of key enzyme namely phosphoketolase in pentose phosphate pathway. Carbohydrate fermentation tests showed AGA58 cannot ferment pentoses which was also confirmed in silico. Putative pyruvate metabolism revealed formate, malate, oxaloacetate, acetate, acetaldehyde, acetoin and lactate forms from pyruvate. AGA58 predicted to carry bacteriocin genes for type A2 lantipeptide, Blp family class II bacteriocins showing antimicrobial potential of this bacterium which can be linked to antagonism tests that AGA58 can inhibit *E. coli* O157:H7, *S. Typhimurium* ATCC14028, and *K. pneumonia* ATCC13883. Moreoever, AGA58 is tolerant to acid and bile concentrations simulating the human gastrointestinal conditions. *L. nagelii* AGA58 depicting the probiotic potential of AGA58 as a first report in literature within same species.

## Introduction

Lactobacilli species are rod-shaped, gram-positive, facultative anaerobe, catalase-negative microorganisms that are heavily being utilized in the food industry due to their well-established technological features and well-documented beneficial effects on health [1]. The Lactobacilli species are classified as GRAS (Generally Recognized as Safe) by USDA [2]. In addition to their starter and probiotic actions, they are potential bioprotective cultures because of their capability of producing antimicrobials such as nisin, enterocin, pediocin, salivaricin, plantaricin etc. [3], which exist in many fermented dairy and vegetable products [4–6].

The *Lactobacillus* genus is a major and comprehensive group in lactic acid bacteria and was isolated from various ecological niches [2]. Shalgam is a vegetable based lactic acid fermented non-alcoholic beverage specific to Southern Anatolia region known to be a rich source of Lactobacilli species such as *L. plantarum, L. paracasei, L. brevis, L. parabrevis, L. acidophilus, L. gasseri, Lb helveticus, L. reuteri* with the former two species are being predominant organisms [7–9]. Till now, no studies performed on uncovering the Shalgam microbiome have been reported the presence of *Liquorilactobacillus* species such as *Liquorilactobacillus nagelii. Liquorilactobacillus* a lactobacillus from liquids, indicating the isolation of species from liquid environments, including plant sap, water and alcoholic beverages. Generally, it shows a homofermentative lifestyle with carrying mol % GC content of 33.9-40.0 [10, 11]. Except *for L. cacaonum, L. hordei, L. mali*, most of the species are motile. Before re-taxonomic structuring of the *Lactobacillus* genus into 23 new genera, *Liquorilactobacillus* species were regarded as part of the *L. salivarius* group [12]. *L. nagelii* is microaerophilic, mesophilic and grows in MRS, including 5% (w/v) NaCl at pH 4.5 at room temperature. It uses citrate and malate in the existence of six-carbon sugar such as glucose. *L. nagelii* biosynthesizes an exopolysaccharide named dextran which is being produced from sucrose [13]. Previous studies reported that *L. nagelii* was isolated from water kefir, fermented cassava food, wild cocoa bean fermentation and semi-fermented wine, silage fermentation and Kombucha [10, 14–16].

The source of isolation of a microorganism is one of the primary factors determining the metabolic potential of the strain. For example, *L. casei* that was isolated from silage material has a different carbohydrate utilization pattern compared to *L. casei* isolated from cheese microenvironment. Thus, strain-level identification and sequencing the entire genome of lactobacilli species is critical for understanding the strain adaptations to ecologically different conditions [17]. Looking into NCBI public database apart from *L. nagelii* AGA58, which is reported in the present study, only 2 strains of *L. nagelii* whole genome sequence was available as of Nov 28, 2021. This limits the understanding of microbial community dynamics and microbial interactions in plant-based fermented foods microbiome [9].

Shalgam at onset of lactic acid fermentation contains sucrose, glucose and fructose [18, 19]. When the fermentation ceases, a residual amount of sugar exists in the final product meaning fermentable carbohydrates are being utilized by the Shalgam microbiome [20]. The microbiome of Shalgam is made up of lactic acid bacteria and yeasts although most research related to Shalgam is focused on former organisms [7]. Since traditional manufacturing of Shalgam relies on spontaneous fermentation, the microbial community is potentially very diverse in terms of harboring unique lactic acid bacteria strains. We isolated a novel *Liquorilactobacillus nagelii* strain AGA58 (Accession No. JAFFQQ000000000) from fermented Shalgam manufactured in the Southern region of Turkey. The aim of the present study was to explore functional and genomic characteristics of *L. nagelii* AGA58 strain isolated from Shalgam for the first time. The strain’s phenotypic features were investigated by screening acid and bile tolerance, carbohydrate fermentation profiles, resistomes and comparative genome analysis with closely related strains. The whole genome of the *L. nagelii* AGA58 using Next-Generation sequencing and “*in silico*” metabolic potentials using bioinformatic tools have also been explored. This is the first report describing genomic and functional characteristics in addition to *in vitro* probiotic potentials of a novel *L. nagelii* AGA58 strain isolated from a plant-based traditional fermented beverage “Shalgam”.

## Materials and Methods

### Isolation of Bacterial Strain and Growth Conditions

*Liquorilactobacillus nagelii* AGA58 was isolated from fermented turnip juice (Shalgam) produced in Adana, Turkey. A 10 mL of the Shalgam sample was diluted with 90mL of Maximum Recovery Diluent (Merck, GmbH, Darmstadt, Germany) in Schott bottle and vortexed for 1 minute with highspeed vortex (MS-3 Basic, IKA-Werke GmbH, Staufen, Germany). A 100µL of sample from serial dilutions were spread on MRS agar (Merck) followed by incubation at 30LJ for 48h anaerobically. The isolate later named as AGA58 was picked and subjected to colony purification twice. Gram staining and catalase tests were performed to pure isolate of *L. nagelii* AGA58, respectively. The cryovial stocks of AGA58 was prepared using MRS broth (Merck) with 25% glycerol and were stored at −80□.

### Genomic DNA extraction, whole-genome sequencing and *de novo* assembly

*Liquorilactobacillus nagelii* AGA58 cryo-culture was sub-cultured twice in MRS broth and anaerobically incubated for 24h at 30□. A 1 mL fresh culture was transferred to a sterile 2 ml microcentrifuge tube and centrifuged at 6000x g for 10min at 4L□. The supernatant was removed. Total genomic DNA extraction was performed in cell pellet by PureLink Genomic DNA Mini Kit (Invitrogen, Thermo-Fisher Scientific, Carlsbad, CA, USA) per manufacturer’s recommendations. The quality and concentration of genomic DNA were checked by a Qubit 3.0 fluorometer (Invitrogen, Thermo-Fisher Scientific) and agarose gel electrophoresis (1%). The sequencing libraries were constructed using Nextera XT DNA Library Preparation Kit (Illumina, San Diego, CA, USA) and sequencing was conducted by Illumina Novaseq platform as paired-end (PE) 2×250 bases read. The low-quality reads were filtered and assembled in the genome assembly service of PATRIC 3.6.12. (https://https://patricbrc.org/app/Assembly2) using auto strategy [21].

### Bioinformatic analyses

Genome annotation and comprehensive genome analysis were conducted using NCBI Prokaryotic Genome Annotation Pipeline (PGAP) and PATRIC 3.6.12. platform [21, 22]. The calculation of orthologous average nucleotide identity values (OrthoANI) of AGA58 and other *Liquorilactobacillus* strains were performed by OrthoANI tool v0.93.1 [23]. Prediction of metabolic pathways of *L. nagelii* AGA58 was carried out using BlastKOALA for screening against KEGG database [24]. The bacteriocin biosynthesis responsible gene cluster prediction was implemented using the BAGEL 4 webserver (http://bagel4.molgenrug.nl/). Then, each member of predicted gene clusters was confirmed by NCBI protein BLAST suite (https://blast.ncbi.nlm.nih.gov/Blast.cgi). The prophage regions on AGA58 genome were identified and annotated with the PHASTER-Phage Search Tool Enhanced Release [25]. A resistome screening was executed by screening whole genome sequences against ResFinder 4.1, CARD, PATRIC 3.6.8 and KEGG databases [21, 24, 26, 27]. To determine the evolutionary relationship within Lactobacillales family, a phylogenetic tree was created per Makarova et al., (2006) [28]. The 16s rRNA nucleotide sequences for each strain and a multi-alignment of the sequences were processed by the MUSCLE algorithm using MEGA software version X (http://www.megasoftware.net). The whole-genome sequence of *Liquorilactobacillus nagelii* AGA58 has been submitted to NCBI under bioproject number PRJNA697983.

### Carbohydrate fermentation

The carbohydrate fermentation patterns of the AGA58 strain were determined using an API 50 CHL kit (BioMérieux, Marcy l’Etoile, France) consisting of 49 different carbohydrate tests under manufacturer’s protocols.

### Determination of antibiotic susceptibility

Antibiogram tests were performed to screen resistance or sensitivity of AGA58 against commonly used antibiotics. Commercially available antibiotic disks [methicillin, vancomycin, amikacin, kanamycin, azithromycin, tetracycline, penicillin G (Bioanalyse, Yenimahalle, Ankara, Turkey); ampicillin, oxacillin, carbenicillin, amoxicillin, streptomycin, erythromycin, rifampicin (Oxoid, Basingstoke, Hampshire, UK)] were utilized to determine antibiotic susceptibility of *L. nagelii* AGA58. The disk diffusion assay was performed per modified Kirby-Bauer method [29]. Interpretation of inhibition zone (mm) results was performed per Clinical and Laboratory Standards Institute’s performance standards for antimicrobial testing [30]. Inhibition zone less than or equal to 14 mm was noted resistant (R). Inhibition zone greater than 20 mm was regarded sensitive (S). Zones between 15 and 19mm were recorded as semi-sensitive or intermediate (I).

### β-haemolytic Activity Tests

Evaluation of β-haemolytic activity of the AGA58 strain was performed via 5% sheep blood containing Columbia agar plate [31]. The isolate was plated on the Columbia agar and incubated at 37LJ for 48h under anaerobic conditions.

### Antibacterial Activity Test

Antibacterial activity test was performed by the agar well diffusion method according to Mishra&Prasad (2005) [32]. The supernatant of 18-20 h grown AGA58 was tested against *E. coli* O157:H7 (ATCC 43895), *S. aureus* (ATCC 25923), *B. cereus* (ATCC 33019), *S. enterica* sv. *Typhimurium* (ATCC 14028), *P. vulgaris* (ATCC 8427) and *K. pneumoniae* (ATCC 13883).

### Growth Kinetics

Growth kinetics of AGA58 were evaluated at different pH and bile concentrations to determine fundamental probiotic function of this organism. Oxyrase (Sigma-Aldrich, USA) enzyme was used to reduce the oxygen level in a microtiter plate. The pH of the MRS medium was adjusted to 6.8-8.4 which is optimum for oxyrase. Oxyrase was added to the medium in proportions, according to McMahon et al., (2020) [33]. MRS medium with five different pH values (pH 2, 3, 4, 5, 7) was prepared using 3N HCl and 3N NaOH and four different bile concentrations (0% control, 0.3%, 0.5% and 1%) was prepared using ox bile extract (Sigma, Germany) followed by incubation at 36.5°C for 30 minutes to activate oxyrase. Growth measurements were performed in HIDEX Sense Microplate Reader (Hidex, Finland) using 96 well-plates with lid. Each well was inoculated with 200 µl of overnight grown culture incubated at 30°C. Each sample was run in quadruplicates. Spectrophotometric measurements were carried out at 30 and 37°C at 300-rpm orbital shake. The OD_600_ measurement was performed every 20 minutes at 72 hours of post-inoculation.

## Results AND DISCUSSION

### Genomic characterization of L. nagelii AGA58

*L. nagelii* AGA58 is a gram-positive, catalase-negative, rod-shaped, lactobacilli species isolated from a traditional fermented turnip beverage called Shalgam produced in Southern Anatolia region. The genome of AGA58 consists of a single linear chromosome with a genomic size of 2,294,535 bp, 36.9% GC content, 2249 protein CDS, 45 tRNA, 3 rRNA (Fig 1 and Table 1), 40 CRISPR repeat region, 38 CRISPR spacer region and 2 CRISPR array region. The genome harbors 1 intact prophage (Figure S3, three bacteriocins (type A2 lantipeptide, BLP family Class II bacteriocins carrying two subunits) in the same operon (Table S3). The KEGG orthology (KO) functional categories of identified protein-coding sequences in the genome of *Liquorilactobacillus nagelii* AGA58 is shown in Table S8. Majority of genes involved in genetic information processing, carbohydrate fermentation, and amino acid metabolism indicate this organism requires various carbon and nitrogen sources to maintain its lifestyle. AGA58 genome predicted to carry glutamine synthetase Type I [E.C.6.3.1.2] which could potentially biosynthesize L-glutamine from ammonia [34]. As most *Liquorilactobacillus* species *L. nagelii* AGA58 is thought to be motile which is predicted from genome analysis. For example, AGA58 genome carries flagellar biosynthesis proteins *FliP, FliQ, FliR, FlhA, FlhB* and flagellar hook protein *FlgE* (Fig 5) which lead us to believe that this novel strain from fermented turnip is likely a motile organism.

**Table 1.** Genomic characteristics of *Liquorilactobacillus nagelii* AGA 58, DSM 13675 and TMW 1.1827

| **Feature** | **AGA 58** | **DSM 13675** | **TMW 1.1827** |
| --- | --- | --- | --- |
| Size (bp) | 2,294,535 | 2,487,096 | 2,406,166 |
| GC content (%) | 36.9 | 36.8 | 36.7 |
| Genes (total) | 2249 | 2461 | 2321 |
| Protein coding sequences | 2151 | 2330 | 2205 |
| tRNA | 45 | 57 | 57 |
| rRNA | 4 | 18 | 18 |
| Non-coding RNA | 4 | 4 | 4 |
| Pseudogenes | 45 | 52 | 37 |
| Plasmid | None | 1 | 3 |

**Table 3.** Comparison of carbohydrate fermentation patterns of *Liquorilactobacillus nagelii* AGA58 and DSM 13675 strains^1^.

| Sugar | Strain |  |
| --- | --- | --- |
|  | AGA 58<sup>2</sup> | DSM 13675<sup>3</sup> |
| Control | - | - |
| Glycerol | - | - |
| Erythritol | - | - |
| D-Arabinose | - | - |
| L-Arabinose | - | - |
| D-Ribose | - | - |
| D-Xylose | - | - |
| L-Xylose | - | - |
| Adonitol | - | - |
| Methyl-βD-xylopyranoside | - | - |
| D-Galactose | + | + |
| D-Glucose | + | + |
| D-Fructose | + | + |
| D-Mannose | + | + |
| **D-Sorbose** | - | + |
| **D-Rhamnose** | - | + |
| Dulcitol | - | - |
| Inositol | - | - |
| **D-Mannitol** | - | + |
| **D-Sorbitol** | - | + |
| Methyl-αD-mannopyranoside | - | - |
| **Methyl-αD-glucopyranoside** | - | + |
| N-Acetylglucosamine | + | + |
| **Amygdalin** | - | + |
| Arbutin | - | - |
| Esculin ferric citrate | + | + |
| Salicin | + | + |
| D-Cellobiose | + | + |
| D-Maltose | + | + |
| D-Lactose | - | - |
| D-Melibiose | - | - |
| D-Sucrose | + | + |
| D-Trehalose | + | + |
| Inulin | - | - |
| D-Melezitose | - | - |
| D-Raffinose | - | - |
| Amidon (Starch) | - | - |
| Glycogen | - | - |
| Xylitol | - | - |
| Gentiobiose | + | + |
| D-Turanose | - | - |
| D-Lyxose | - | - |
| **D-Tagatose** | - | + |
| D-Fucose | - | - |
| L-Fucose | - | - |
| D-Arabitol | - | - |
| L-Arabitol | - | - |
| Gluconate | - | - |
| 2-Keto-gluconate | - | - |
| 5-Keto-gluconate | - | - |

<sup>1</sup>Fermentation results are expressed as follows; (+) positive, (-) negative.
<sup>2</sup>AGA58 strain has been isolated first in this study.
<sup>3</sup>DSM13675 strain was previously studied by Buron-Moles et. aLb. (2019).

**Fig 1.**
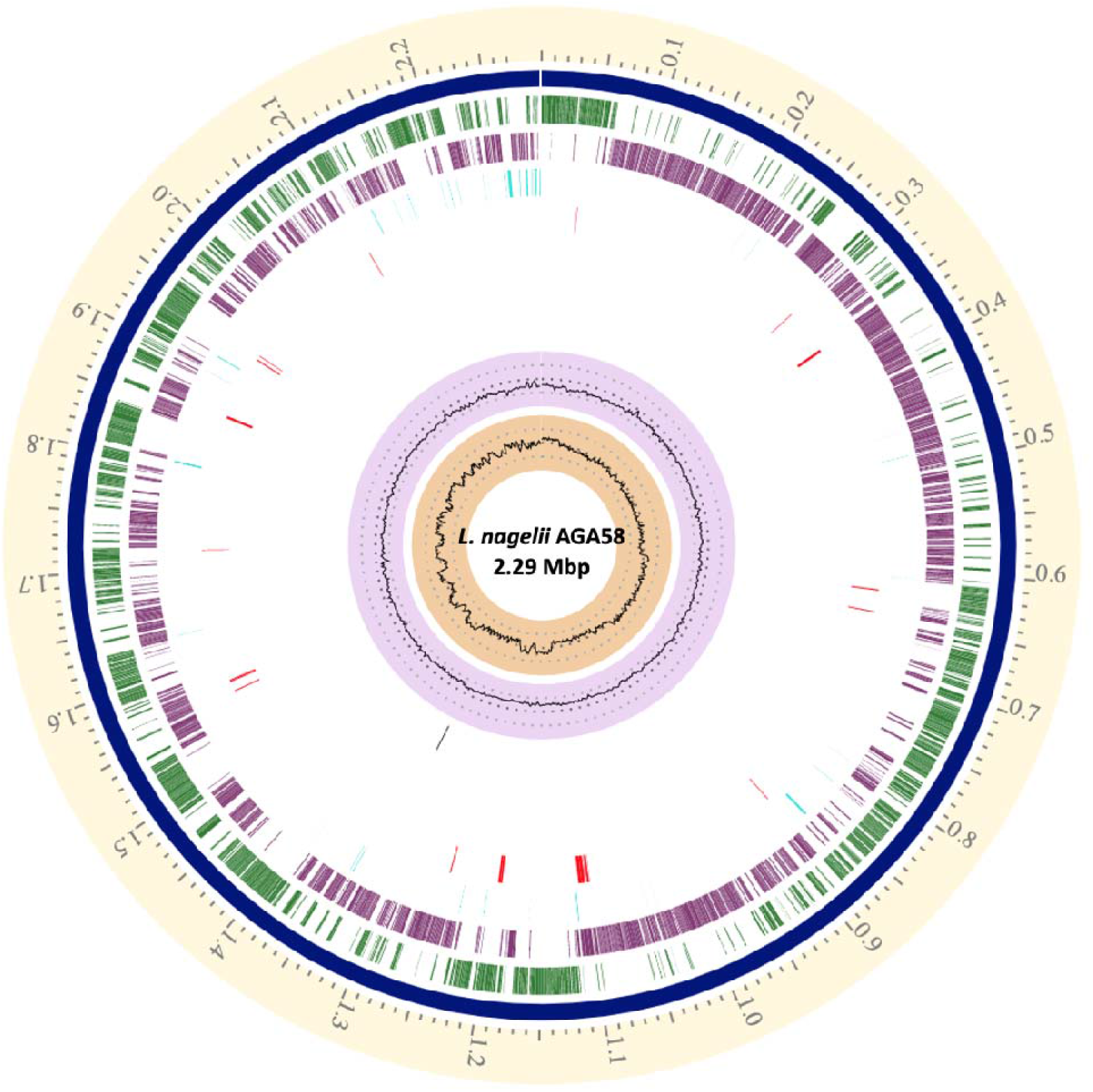
Genome Atlas of *L. nageli* AGA58. Light yellow circle (first circle from outside to inside) shows position label (Mbp). Second outer dark blue circle displays chromosome. Third outer green and forth outer purple circles show forward and reverse coding sequences, respectively. Turquise color displays non codig sequence features and red color displays AMR genes. Lilac coloured second inner circle displays the GC content of the genome. The first inner circle depicts the GC skew (G-C)/(G+C).

### Bacteriocins and Antibiotic Resistome

Bacteriocin screening via BAGEL4 revealed that AGA58 predicted to carry 3 core bacteriocin biosynthesis genes in the same operon (Fig 4). The protein sequence encoding A2 lantipeptide did also show around 80% similarity with Plantaricin_423 peptide which is known to harbor anti-listerial activity [35]. However, performing a protein blast of the sequence [*MKKEIELSEKELVRII****GG****KYYGNGVSCTKKHGCKVNWGQAFTCSVNRFANFGHGNC]* in NCBI showed that it has the highest sequence homology with A2 lantipeptide (Table S2, supplemental). Besides, AGA58 genome predicted to carry 2 other Blp family class II bacteriocins (Fig 3) which is a family of bacteriocidal bacteriocins extracellularly secreted to combat Gram-positive bacteria that are closely related [36]. With AGA58 possessing such bacteriocidal gene cluster allows us to link the genomic evidence of bacteriocin presence with in vitro antibacterial activity test results showing the antagonistic effect of supernate of AGA58 grown media against *E. coli* O157:H7 ATCC 43895, *S. enterica* sv.*Typhimurium* ATCC14028, and *K. pneumonia* ATCC13883.

In vitro antibiotic sensitivity test results revealed that AGA58 is sensitive to ampicillin (10 ug), carbonicillin (100ug), erythromycin (10 ug), penicillin G (10 U), tetracycline (30mcg) rifampicin (5 ug) amoxycillin (25 ug) with the following zones of inhibition achieved: 11.27 mm, 15.75 mm, 18.55 mm, 21.25 mm, 15.53 mm, 25.78 mm, 24.12 mm, 17.71 mm, respectively. On the other hand, azithromycin, oxacillin, streptomycin, vancomycin, methicillin, amikacin, kanamycin provided lower than 15 mm inhibition zones which shows AGA58 is resistant to those antibiotics. Looking into the genome, however, only vancomycin and penicillin resistance genes were found in AGA58, which reveals that phenotype and genotype does not overlap completely [37].

### Comparative genome analysis of L. nagelii AGA58

Based on orthoANI and 16s rRNA sequences alignments, *L. nagelii* AGA58 is closely related to other *L. nagelii* strains of TMW 1.1827 and DSM 13675, which were being isolated from water kefir and semi fermented wine, respectively (Fig 2 and 3). Next closely related neighbors found were *L. hordei* TMW1.1822 and *L. mali* LM596 which were being isolated from water kefir and apple juice, respectively. *L. nagelii* AGA58 carries a single intact prophage, whereas *L. nagelii* TMW 1.1827 possesses no intact prophage though another strain of *L. nagelii* DSM 13675 harbours 2 intact prophages. *L. nagelii* TMW 1.1827, whose genome sequence is publicly available in NCBI possesses a genomic size of 2.41 Mbp and shows a GC content of 36.68% and a total number of 2,391 CDS, including all three plasmids (Table 1). Until 2015, the only publicly available whole-genome sequences of *L. nagelii* strains yielded from a comparative genomics project together against 211 other lactic acid bacteria [38]. *L. nagelii* AGA58 was isolated from the fermented turnip microenvironment and associated with a different ecosystem than wine thus coming across with different conditions. Those differences in the adaptation to distinct microenvironments were also evident in the genomes. For example, the annotated differences between AGA58 and DSM 13675 strains from a fermented turnip, semi fermented wine could be explained by genes associated with carbohydrate metabolism in particular enzymes of citrate and accompanying acetolactate metabolism. These enzyme encoding genes only existed in shalgam and water kefir isolate TMW 1.1827. The AGA58 is distinguished from DSM13675, TMW 1.1827 with galactose metabolism with AGA58 using Leloir pathway to uptake galactose by specific galactose permeases versus latter two utilize Tagatose shunt (Table S1, S2, Fig 6). This is also evident in API 50 CHL carbohydrate fermentations test that AGA58 did not ferment D-Tagatose although DSM13675 did (Table 2) Since galactose sugar is not readily available in respective environments, a challenge remains in explaining specific adaptations based on isolation sources. The genomic evidence predicting environmental adaptations seen in malted barley isolate *L. hordeii* DSM 19519 [38] or water kefir isolate *L. hordeii* TMW 1.1822 [39, 40] is more pronounced and decisive in sucrose metabolism. Comparative analysis of whole-genome sequences of TMW 1.1822 vs TMW 1.1827 revealed core genome of two strains contains 1380 CDS representing 56% and 57.7% of the strains, respectively. The major components of core genes were associated with protein, carbohydrate and amino acid metabolism. However, TMW 1.1822 accessory genome predominantly carries additional genes for amino acid and carbohydrate metabolism plus cell wall biosynthesis as opposed to TMW 1.1827. Because both TMW 1.1822 and TMW 1.1827 are related to the sugar-rich water kefir environment, it was observed that both strains are adapted to this condition by supplementary genes encoding carbohydrate metabolism. Since *L. nagelii* strains do not generally exist in sourdoughs, they cannot proliferate during sourdough fermentation. This might be due to strain adaptations to microenvironments they were being originally isolated (shalgam, partially fermented wine, water kefir) which are not substitutable with the sourdough matrice. The fact that the shalgam ecosystem harbours diverse lactic acid bacteria species (in addition to *L. plantarum, L. paracasei, L. brevis, L. parabrevis, L. acidophilus, L. gasseri, Lb helveticus, L. reuteri)* [7–9] and yeast species reveals this is a unique complex fermented ecological system[20].

**Fig 2.**
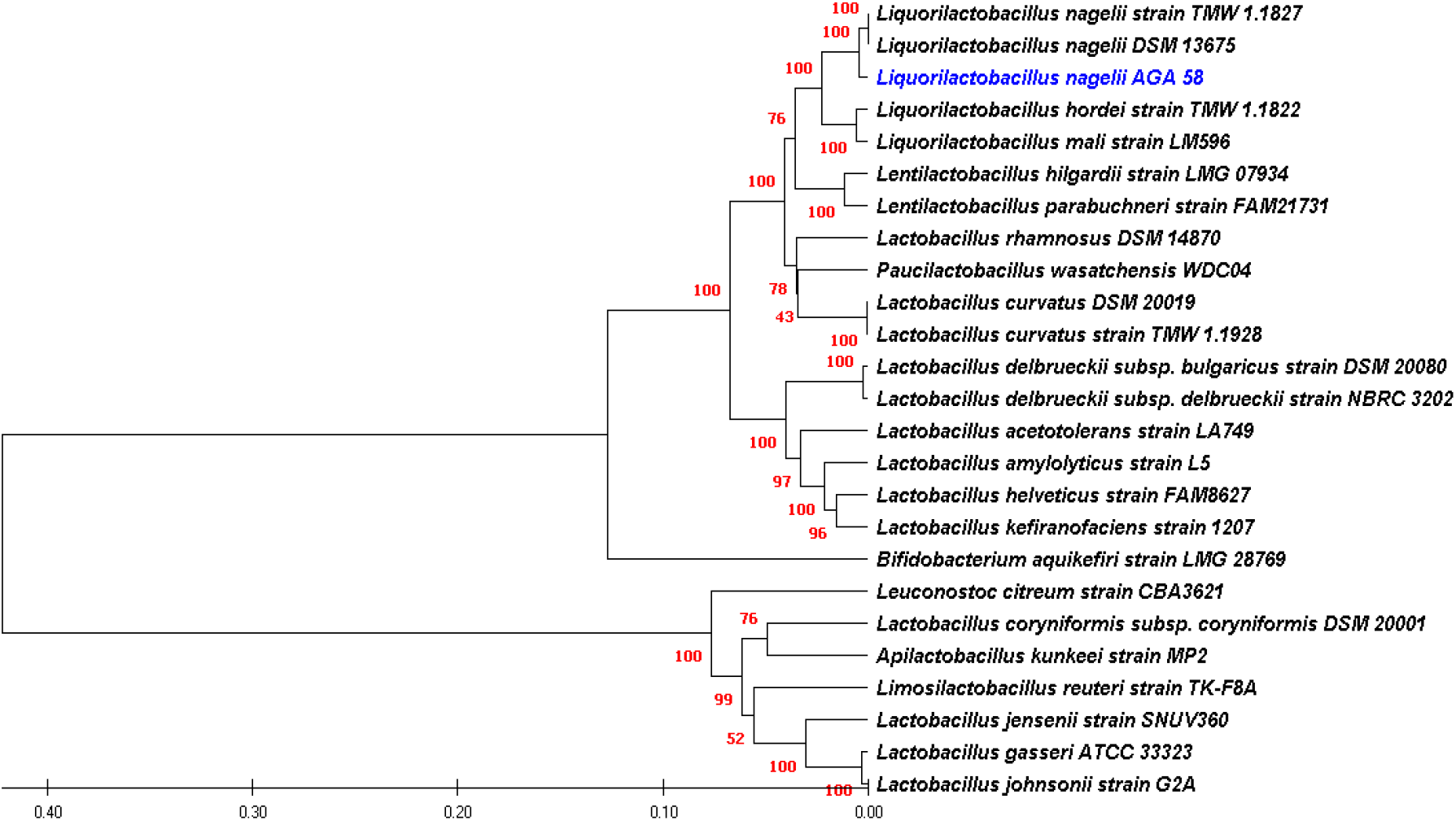
Phylogenetic tree of *L. nagelii AGA58* versus other lactic acid bacteria species and *Bifidobacterium aquikefiri* based on 16s rRNA sequences

**Fig 3.**
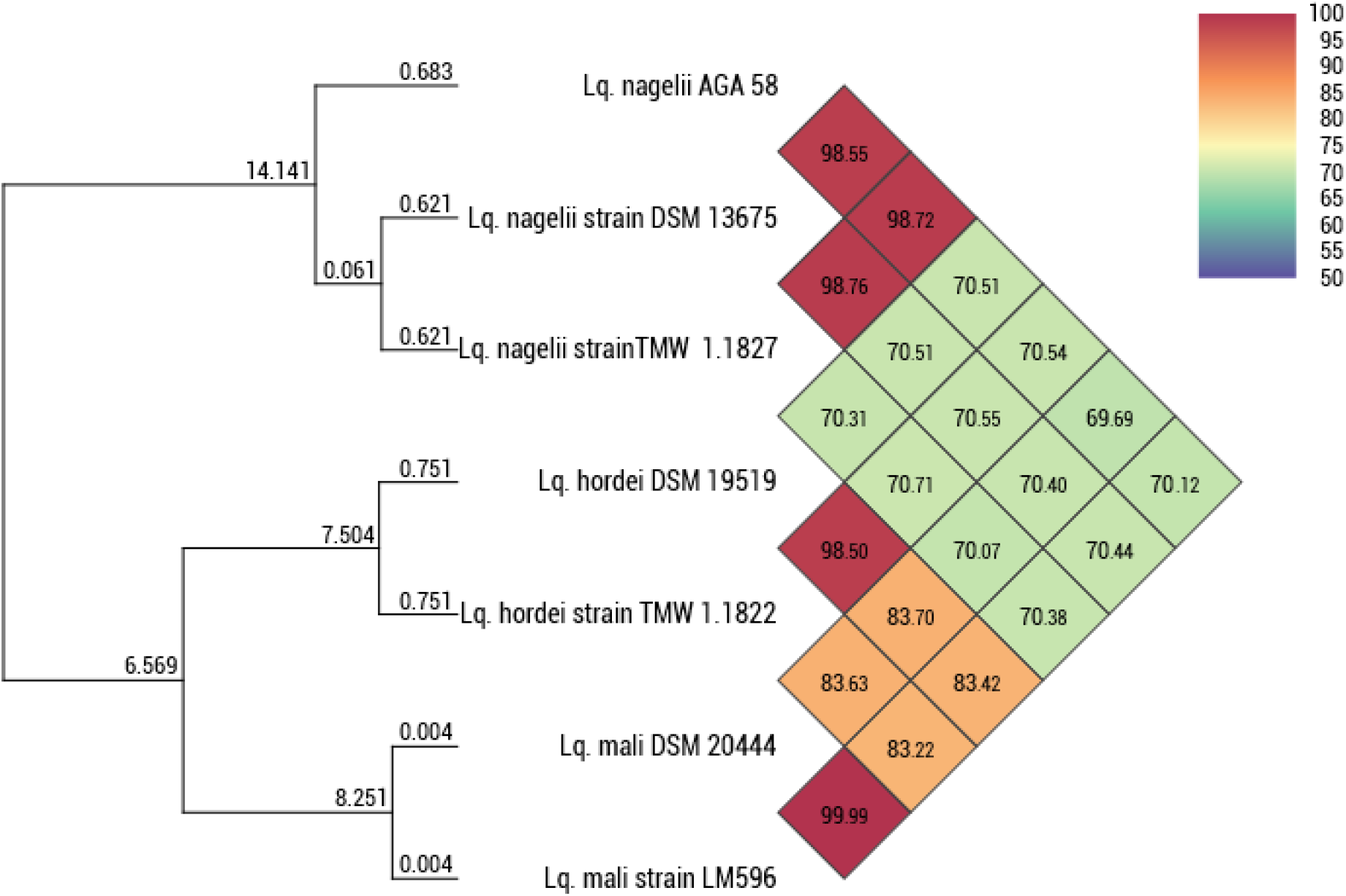
Orthologous Average Nucleotide Identity dendogram between *L. nagelii* AGA58 vs other *Liquorilactobacillus* species.

**Fig 4.**
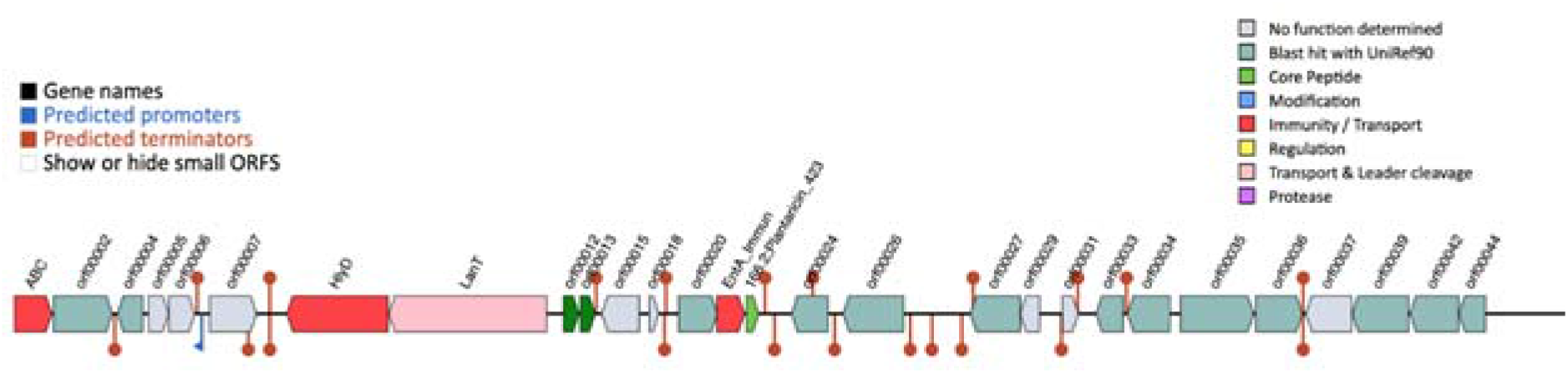
The predicted gene cluster responsible for the biosynthesis of bacteriocins by using BAGEL4 webserver. The dark green genes orf00012 and 00013 encodes blp II class bacteriocins, and Light green 166:2; Plantaricin_423 gene encodes Type A2 lantipeptide per NCBI-protein blast analysis.

**Fig 5.**
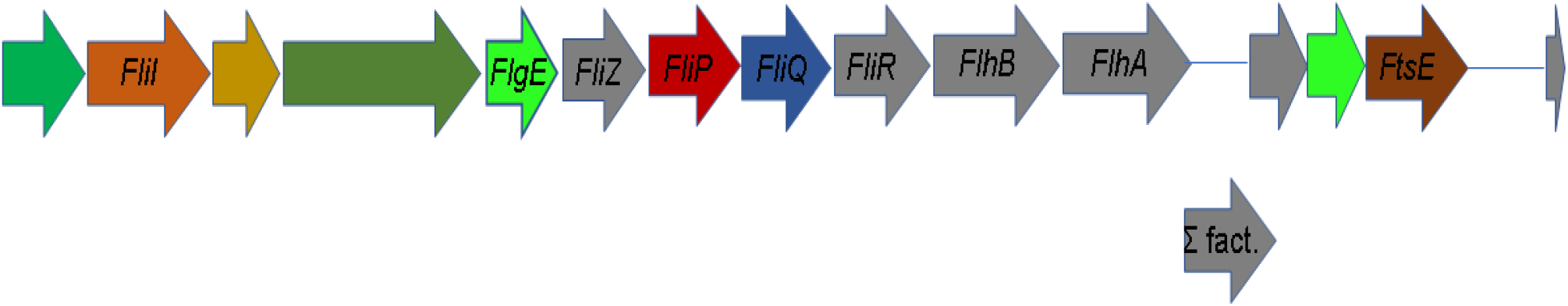
Predicted flagellar biosynthesis enzyme encoding gene cluster (∑ fact: Sigma factor for flagellar gene cluster)

**Fig 6.**
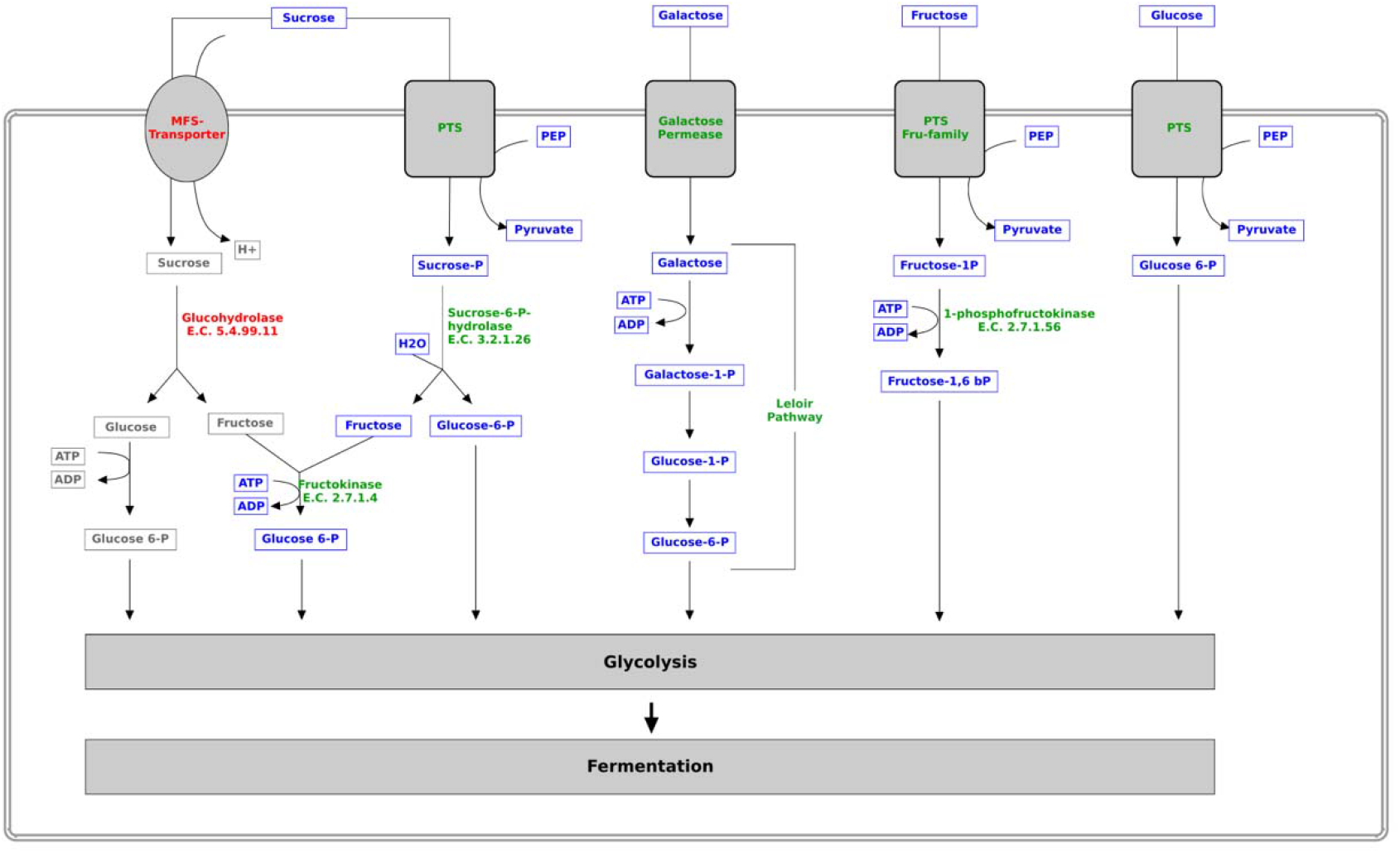
Putative carbohydrate metabolism overview of *L. nagelii* AGA58 per genomic analysis. Green font shows the presence of the enzyme, red font depicts the absence of enzyme in the genome of AGA58.

In addition, lactic acid bacteria commonly exist in cereal fermentations are usually associated with animals (e.g., gut microbiota; *L. fermentum, L. reuteri, L. sakei*) or vegetable fermentations (e.g., *L. plantarum*) [41]. Although *L. nagelii* isolates were mainly related to alcoholic fermentations such as stuck wine and water kefir which perhaps shows the alcohol tolerance of this species, it can also exist other fermented plant material related non-alcoholic conditions [10, 42, 43]. *L. nagelii* is known for its dextran biosynthesis from sucrose via dextran sucrase enzyme [10], however, AGA58 genome carries a similar protein sequence that provides a significant E-value upon protein blast though the gene is annotated as hypothetical protein in RAST (data not shown) (https://rast.nmpdr.org).

### Carbohydrate Metabolism

The genomic analysis and in vitro carbohydrate fermentation profiles of *L. nagelii* AGA58 revealed this strain is capable of metabolizing galactose, glucose, fructose, sucrose, mannose, N-acetyl glucosamine, maltose, trehalose, esculin, ferric citrate, salicin, D-cellobiose, D-maltose, Gentiobiose sugars (Table 2). For example, dextrose, one of the easier carbon source which is mainly utilized in MRS medium, fluxes into the glycolysis pathway when *L. nagelii* AGA58 is grown with this 6-carbon sugar (Fig 6). Fig S1 and Fig S2 show the genome predicted carbohydrate metabolism pathways for *L. nagelii* AGA58. For example, D-glucose molecules in the medium enter the cytoplasm thru PTS and they were being phosphorylated into Glucose-6P followed by conversion into Fructose 1,6 biphosphate via *pfk* gene encoding phosphofructokinase (Fig 6). Later, it feeds into the lower half of the glycolysis pathway by *fba* gene encoding fructose biphosphate aldolase. Since AGA58 possesses those key enzymes for the glycolytic pathway, this strain ferments the abovementioned six-carbon sugars to 2 moles ATP and 2 moles of lactate per each mole of sugar consumed (Table S1, Table S2) [44].

Fructose, another hexose sugar available in the shalgam microenvironment is also metabolized by AGA58 (Table 2, Fig 6). D-Fructose is first phosphorylated to beta-D-Fructose-6P by *fk* gene encoding fructokinase, which further phosphorylated to beta-D-Fructose-1,6 P2 followed by being converted to Glceraldehyde-3P via *fba* gene encoding fructose biphosphate aldolase enzyme. Eventually, D-fructose is being metabolized thru the glycolysis shunt and generates both ATP and lactate as a result of homofermentative lifestyle (Table S1, Table S2) [44].

Genome analysis of *L. nagelii* AGA58 revealed that galactose is fermented through Leloir pathway where it enters the cell via galactose specific permease after which it is phosphorylated to Galactose-1-P at the expense of 1 ATP followed by transforming to Glucose-1-P then Glucose-6-P which feeds into the glycolysis shunt resulting in lactate biosynthesis (Fig 6) [44]. Compared to TMW 1.1827 and DSM 13675 which use the Tagatose path for galactose metabolism with DSM 13675 fermenting D-tagatose (Table 2), AGA58 prefers Leloir route (Fig 6) as no acid production from Tagatose observed in API panel (Table 2). It is worth mentioning here all three *L. nagelii* strains are capable of fermenting sucrose which is one of the primary carbon source available in all three ecosystems at the onset of lactic acid fermentations. However, main difference in sucrose metabolism which is evident from functional comparative genome analysis output is that TMW 1.1827 and DSM 13675 harbor sucrose permease for sugar uptake versus AGA58 is predicted to carry PTS type sucrose specific transport system, which further flows to glycolysis shunt upon cleavage by sucrase to glucose and fructose moieties (Table 2, Fig S1, S2).

DSM 13675 carries transketolase but AGA58 does not. Transketolase is functional in condensing pentose sugars into hexoses so that strains can build cell wall material when the only fermentable sugar is 5-carbon one in respective environments [45]. Thus, it is predicted that AGA58 perhaps is not capable of biosynthesizing cell wall material from pentose sugars rather they are being utilized in purine and pyrimidine synthesis (Fig S1 and S2) which is in alignment with API 50 CHL results that AGA58 did not produce acid from any of the pentose sugars provided in Table 2.

### Pyruvate Metabolism

Pyruvate is a critical metabolite in many fermentations for serving as an electron acceptor for NADH to NAD+ regeneration step to maintain redox balance. Under certain conditions, lactic acid bacteria utilize alternate pathways for using pyruvate than the conversion to lactate. The alternative ways in which *L. nagelii* AGA58 predicted to utilize pyruvate is shown in Fig 7. Putative pyruvate metabolism of AGA58 revealed formate, malate, oxaloacetate, acetate, acetaldehyde, acetoin and lactate could form from pyruvate. Some of these reactions could occur even regular glucose fermentation and might serve an anabolic role for example production of acetyl CoA could be necessary for lipid biosynthesis (Fig. 7) [44]. Genomic analysis indicated existence of enzymes participating in different downstream pyruvate transformation. AGA58 is not only capable of ATP generation upon acetate formation by an acetate kinase, but also for maintenance of redox balance by L/D-lactate (Fig. 7, Table S6). Since no alcohol dehydrogenase (*adh*) enzyme encoding gene was found in AGA58 genome, other strategies for NAD+ re-cycling could be more favorable such as transformation of pyruvate to acetoin. For catabolism of pyruvate to α-acetolactate, acetolactate synthase is the key enzyme found in AGA58 genome for synthesis of these compounds. The α-acetolactate is not stable a molecule thus it is reduced to acetoin by α-acetolactate decarboxylase or decarboxylated to diacetyl non-enzymatically. Finally, diacetyl/acetoin reductase enzyme reduces diacetyl to acetoin, which is then transformed to 2,3-butanediol upon concurrent NAD+ regeneration [40]. Since AGA58 was found to carry all relevant enzyme encoding genes in its genome for the biosynthesis of acetoin, acetaldehyde and acetate it might contribute to flavor profile of Shalgam (Table S6). Although carries a citrate synthase, AGA58 lacks the remaining key enzymes completing the TCA cycle which is in alignment with *L. hordei* TMW 1.1822 isolated from water kefir [40].

**Fig 7.**
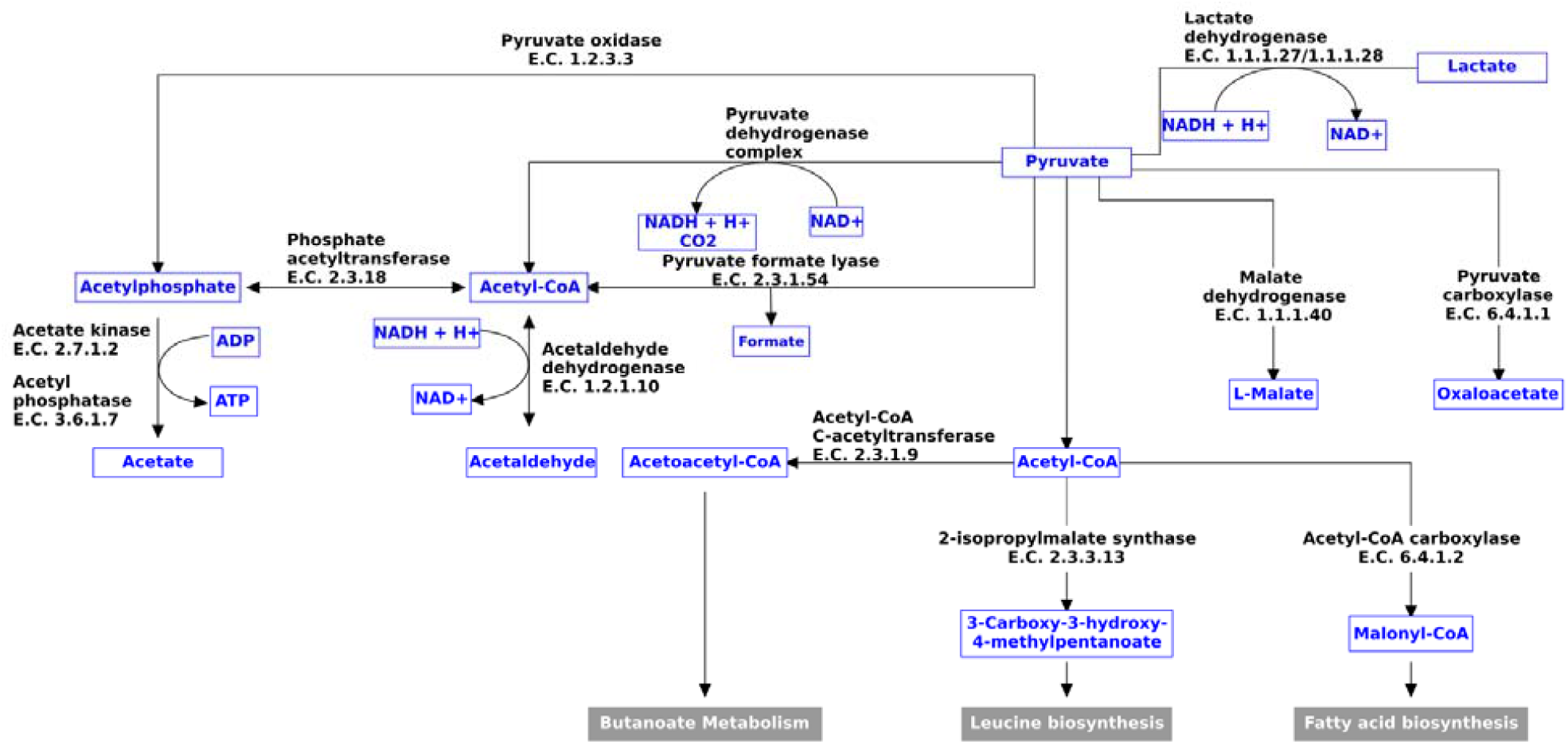
Putative pyruvate metabolism of *L. nagelii* AGA58 revealed from genome analysis.

### Acid and Bile Tolerance

Acid challenge tests of AGA58 performed at pH= 2, 3, 4, 5, 7 are shown as growth curves in Fig S6. Obviously, no or very poor growth is seen at pH=2 and pH=3. However, AGA58 growth rate gradually increased when it was grown in pH=4, pH=5 or pH=7, respectively. The maximum specific growth rates (µmax) achieved were pH=7>pH=5>pH=4. Although pH=7 provided faster µmax compared to pH=5 condition, final biomass concentrations achieved were similar. On the other hand, pH=4 condition achieved final biomass concentration was remarkably lower compared to both pH=5 and pH=7. This perhaps relates to the fact that cells tend to maintain neutral pH even when placed into harsh acidic conditions that requires ATP facilitated export of H+ cations across the cytoplasm [44]. Expenditure of ATP towards stress response would limit the overall growth rate of the strain as the cellular machinery is directed to both cell growth and acid stress combat. For pH=5, the balance of stress response and cell growth only impacted the umax and perhaps did not exceed the threshold thus, AGA58 could still grow to similar cell densities with pH=7 neutral condition. However, when AGA58 was grown on pH=4, both cell growth rate and final cell density were impacted, perhaps owing to the cell combatting for higher acid stress still attempting to increase the biomass where the threshold probably exceeded; thus, overall cell biomass had a lower yield.

Bile salt challenge tests performed across 0.3%, 0.5%, 1% bile salt treatments provided no significantly different doubling times for AGA58 (P>0.05). Control treatment (i.e. no bile) yielded a significantly shorter doubling time than all the others (P<0.05). Similarly, umax values obtained across all bile challenges were not significantly different. Again, control treatment revealed the highest maximum specific growth rate. A slowdown in doubling time and umax might relate to the absence of *bshA* gene encoding bile salt hydrolase in AGA58 genome. On the contrary, a considerable amount of biomass growth achieved even at 1% bile salt treatment indicates that AGA58 persists resilience to bile concentration at or above seen in human gut conditions. Similar results were also reported for another Shalgam microbiome member L. plantarum DY46 which showed a slowdown in growth rate and in the presence of acid or bile salt conditions [9].

*Liquorilactobacillus nagelii* AGA58, like most other lactic acid bacteria found in fermented foods microbiome, is a facultative anaerobic bacterium that will grow best at zero or very low oxygen levels. Because the surface area to volume ratio of 200 µl of liquid in 96-well plates is relatively high compared with test tubes, the dissolved oxygen in the medium slows its growth [33]. By utilizing Oxyrase enzyme to achieve a micro-anaerobic environment by chemically removing oxygen [33] increased the growth rate and reduced the doubling time of AGA58 compared to control (i.e. no oxyrase added). Similar results were also reported by [33] that a oxyrase supplementation to each well of 48 well-plate increased both the rate and extent of growth for a Cheddar cheese isolate *Paulactobacillus wasatchensis* WDC04 [33].

## Conclusion

The putative functional genome analysis and in vitro tests revealed *L. nagaelii* AGA58 harbor the genes associated with sucrose, glucose and fructose metabolism in which those sugars are readily available in Shalgam microenvironment. We found that AGA58 is micro-anaerobic organism predicted to has flagella in alignment with motile character. It is capable of utilizing hexose sugars via glycolysis due to its obligatory homofermentative nature which is also confirmed by genomic evidence that AGA58 possesses the genes encoding phosphofructokinase and fructose biphosphate aldolase. No acid production from pentose sugars observed perhaps relates to the fact that AGA58 does not contain key enzyme phosphoketolase for pentose phosphate pathway; rather, the five-carbon sugars are being converted to purines and pyrimidines, which are building blocks of DNA molecule. Genome analysis predicted galactose is fermented thru Leloir pathway via galactose specific permease which further feeds into glycolysis unlike the water kefir isolate of *L. nagelii* TMW 1.1827 and DSM 13675. Putatve pyruvate pathway revealed formate, acetate, malate, lactate, acetaldehyte and acetoin can be formed from pyruvate. A single intact prophage and transposases found in AGA58 is an indicator of the plasticity of the genome. Predicted novel bacteriocins in the same operon reveal this strain is potentially biosynthesizing antimicrobial peptides, which could be associated with the antibiogram tests showing inhibition zones against *E. coli* O157:H7 ATCC 43895, *S. enterica* sv. *Typhimurium* ATCC 14028 and *K. pneumonia* ATCC13883. Acid and bile tolerance tests showed AGA58 could survive against human simulated gastrointestinal conditions. As a result,

*L. nagelii* AGA58 is a novel homofermentative lactic acid bacterium isolated fermented turnip that carries potential probiotic functional characteristics that require further attention for analyzing probiotic characteristics via in vitro and in vivo tests.

## Acknowledgement

This work has been supported by Erciyes University Scientific Research Projects Unit with grant number FKB-2020-10551. We thank Mr. Ismail Gumustop with excellent support during drawing of metabolic pathways.

## Notes

### Competing Interest Statement

The authors have declared no competing interest.

