## Supplementary Information for "Metabolic potentials of *Liquorilactobacillus nagelii* AGA58 isolated from Shalgam based on genomic and functional analysis"

**Table S1A.** Functional comparison of differences between the strains AGA58 vs DSM 13675 (A=AGA58)

| **Presence** | **Category** | **Subcategory** | **Subsystem** | **Role** | **SS active A** | **SS active B** |
| --- | --- | --- | --- | --- | --- | --- |
| **A** | Amino Acids and Derivatives | Aromatic amino acids and derivatives | Chorismate: Intermediate for synthesis of Tryptophan, PAPA antibiotics, PABA, 3-hydroxyanthranilate and more. | Anthranilate synthase, aminase component (EC 4.1.3.27) | yes | no |
| **A** | Amino Acids and Derivatives | Aromatic amino acids and derivatives | Chorismate: Intermediate for synthesis of Tryptophan, PAPA antibiotics, PABA, 3-hydroxyanthranilate and more. | Indole-3-glycerol phosphate synthase (EC 4.1.1.48) | yes | no |
| **A** | Clustering-based subsystems | May be related to ADP-phosphoribose and NAD-dependent acetylation | CBSS-216591.1.peg.168 | Histone acetyltransferase HPA2 and related acetyltransferases | yes | no |
| **A** | Protein Metabolism | Protein biosynthesis | tRNA aminoacylation, Pro | tRNA proofreading protein STM4549 | yes | no |
| **A** | Secondary Metabolism | Plant Hormones | Auxin biosynthesis | Anthranilate phosphoribosyltransferase (EC 2.4.2.18) | yes | no |
| **A** | Secondary Metabolism | Plant Hormones | Auxin biosynthesis | Phosphoribosylanthranilate isomerase (EC 5.3.1.24) | yes | no |
| **A** | Stress Response | no subcategory | Dimethylarginine metabolism | NG,NG-dimethylarginine dimethylaminohydrolase 1 (EC 3.5.3.18) | yes | no |
| **A** | Virulence, Disease and Defense | Resistance to antibiotics and toxic compounds | Cobalt-zinc-cadmium resistance | Probable cadmium-transporting ATPase (EC 3.6.3.3) | yes | no |
| **A** | Virulence, Disease and Defense | Resistance to antibiotics and toxic compounds | Copper homeostasis | Copper chaperone | yes | no |

**Table S1B.** Functional comparison of differences between the strains AGA58 vs DSM 13675 (B= DSM 13675)

| **Presence** | **Category** | **Subcategory** | **Subsystem** | **Role** | **SS active A** | **SS active B** |
| --- | --- | --- | --- | --- | --- | --- |
| **B** | Amino Acids and Derivatives | Aromatic amino acids and derivatives | Common Pathway For Synthesis of Aromatic Compounds (DAHP synthase to chorismate) | 2-keto-3-deoxy-D-arabino-heptulosonate-7-phosphate synthase I beta (EC 2.5.1.54) | no | yes |
| **B** | Amino Acids and Derivatives | Aromatic amino acids and derivatives | Common Pathway For Synthesis of Aromatic Compounds (DAHP synthase to chorismate) | Shikimate/quinate 5-dehydrogenase I beta (EC 1.1.1.282) | no | yes |
| **B** | Carbohydrates | Central carbohydrate metabolism | Pentose phosphate pathway | Transketolase (EC 2.2.1.1) | no | yes |
| **B** | Carbohydrates | Di- and oligosaccharides | Lactose and Galactose Uptake and Utilization | Galactose-6-phosphate isomerase, LacA subunit (EC 5.3.1.26) | no | yes |
| **B** | Carbohydrates | Di- and oligosaccharides | Lactose and Galactose Uptake and Utilization | Galactose-6-phosphate isomerase, LacB subunit (EC 5.3.1.26) | no | yes |
| **B** | Carbohydrates | Di- and oligosaccharides | Lactose and Galactose Uptake and Utilization | Lactose phosphotransferase system repressor | no | yes |
| **B** | Carbohydrates | Di- and oligosaccharides | Lactose and Galactose Uptake and Utilization | Tagatose-6-phosphate kinase (EC 2.7.1.144) | no | yes |
| **B** | Carbohydrates | Di- and oligosaccharides | Sucrose utilization | Sucrose permease, major facilitator superfamily | no | yes |
| **B** | Carbohydrates | Monosaccharides | D-Galacturonate and D-Glucuronate Utilization | Pectate lyase precursor (EC 4.2.2.2) | no | yes |
| **B** | DNA Metabolism | CRISPs | CRISPRs | CRISPR-associated helicase Cas3 | no | yes |
| **B** | DNA Metabolism | DNA repair | DNA repair, bacterial MutL-MutS system | MutS domain protein, family 4 | no | yes |
| **B** | DNA Metabolism | no subcategory | Restriction-Modification System | Type I restriction-modification system, DNA-methyltransferase subunit M (EC 2.1.1.72) | no | yes |
| **B** | DNA Metabolism | no subcategory | Restriction-Modification System | Type I restriction-modification system, restriction subunit R (EC 3.1.21.3) | no | yes |
| **B** | Phages, Prophages, Transposable elements, Plasmids | Phages, Prophages | Phage capsid proteins | Phage head maturation protease | no | yes |
| **B** | Phages, Prophages, Transposable elements, Plasmids | Phages, Prophages | Phage packaging machinery | Phage portal protein | no | yes |
| **B** | Phages, Prophages, Transposable elements, Plasmids | Phages, Prophages | Phage packaging machinery | Phage terminase, small subunit | no | yes |
| **B** | Regulation and Cell signaling | Programmed Cell Death and Toxin-antitoxin Systems | Toxin-antitoxin replicon stabilization systems | HigB toxin protein | no | yes |
| **B** | Regulation and Cell signaling | Programmed Cell Death and Toxin-antitoxin Systems | Toxin-antitoxin replicon stabilization systems | YoeB toxin protein | no | yes |
| **B** | Respiration | Electron donating reactions | Respiratory dehydrogenases 1 | Glycerol dehydrogenase (EC 1.1.1.6) | no | yes |
| **B** | Virulence, Disease and Defense | Resistance to antibiotics and toxic compounds | Beta-lactamase | Beta-lactamase class A | no | yes |

**Table S2A.** Functional comparison of differences between the strains AGA58 vs TMW 1.1827 (A=AGA58)

| **Presence** | **Category** | **Subcategory** | **Subsystem** | **Role** | **SS active A** | **SS active B** |
| --- | --- | --- | --- | --- | --- | --- |
| **A** | Amino Acids and Derivatives | Aromatic amino acids and derivatives | Chorismate: Intermediate for synthesis of Tryptophan, PAPA antibiotics, PABA, 3-hydroxyanthranilate and more. | Anthranilate synthase, aminase component (EC 4.1.3.27) | yes | no |
| **A** | Amino Acids and Derivatives | Aromatic amino acids and derivatives | Chorismate: Intermediate for synthesis of Tryptophan, PAPA antibiotics, PABA, 3-hydroxyanthranilate and more. | Indole-3-glycerol phosphate synthase (EC 4.1.1.48) | yes | no |
| **A** | Carbohydrates | Aminosugars | Chitin and N-acetylglucosamine utilization | Beta-hexosaminidase (EC 3.2.1.52) | yes | no |
| **A** | Clustering-based subsystems | May be related to ADP-phosphoribose and NAD-dependent acetylation | CBSS-216591.1.peg.168 | Histone acetyltransferase HPA2 and related acetyltransferases | yes | no |
| **A** | Membrane Transport | ABC transporters | ABC transporter oligopeptide (TC 3.A.1.5.1) | Oligopeptide ABC transporter, periplasmic oligopeptide-binding protein OppA (TC 3.A.1.5.1) | yes | no |
| **A** | Phages, Prophages, Transposable elements, Plasmids | Phages, Prophages | Phage capsid proteins | Phage capsid and scaffold | yes | no |
| **A** | Phages, Prophages, Transposable elements, Plasmids | Phages, Prophages | Phage packaging machinery | Phage terminase, large subunit | yes | no |
| **A** | Phosphorus Metabolism | no subcategory | High affinity phosphate transporter and control of PHO regulon | Phosphate regulon transcriptional regulatory protein PhoB (SphR) | yes | no |
| **A** | Secondary Metabolism | Plant Hormones | Auxin biosynthesis | Anthranilate phosphoribosyltransferase (EC 2.4.2.18) | yes | no |
| **A** | Secondary Metabolism | Plant Hormones | Auxin biosynthesis | Phosphoribosylanthranilate isomerase (EC 5.3.1.24) | yes | no |
| **A** | Stress Response | Oxidative stress | Glutathione: Biosynthesis and gamma-glutamyl cycle | Glutathione biosynthesis bifunctional protein gshF (EC 6.3.2.2)(EC 6.3.2.3) | yes | no |
| **A** | Virulence, Disease and Defense | Resistance to antibiotics and toxic compounds | Cobalt-zinc-cadmium resistance | Probable cadmium-transporting ATPase (EC 3.6.3.3) | yes | no |
| **A** | Virulence, Disease and Defense | Resistance to antibiotics and toxic compounds | Copper homeostasis | Copper chaperone | yes | no |

**Table S2B.** Functional comparison of differences between the strains TMW 1.1827 vs AGA58 (B=TMW 1.1827)

| **Presence** | **Category** | **Subcategory** | **Subsystem** | **Role** | **SS active A** | **SS active B** |
| --- | --- | --- | --- | --- | --- | --- |
| B | Amino Acids and Derivatives | Aromatic amino acids and derivatives | Common Pathway For Synthesis of Aromatic Compounds (DAHP synthase to chorismate) | 2-keto-3-deoxy-D-arabino-heptulosonate-7-phosphate synthase I beta (EC 2.5.1.54) | no | yes |
| B | Amino Acids and Derivatives | Aromatic amino acids and derivatives | Common Pathway For Synthesis of Aromatic Compounds (DAHP synthase to chorismate) | Shikimate/quinate 5-dehydrogenase I beta (EC 1.1.1.282) | no | yes |
| B | Carbohydrates | Central carbohydrate metabolism | Pentose phosphate pathway | Transketolase (EC 2.2.1.1) | no | yes |
| B | Carbohydrates | Di- and oligosaccharides | Lactose and Galactose Uptake and Utilization | Galactose-6-phosphate isomerase, LacA subunit (EC 5.3.1.26) | no | yes |
| B | Carbohydrates | Di- and oligosaccharides | Lactose and Galactose Uptake and Utilization | Galactose-6-phosphate isomerase, LacB subunit (EC 5.3.1.26) | no | yes |
| B | Carbohydrates | Di- and oligosaccharides | Lactose and Galactose Uptake and Utilization | Lactose phosphotransferase system repressor | no | yes |
| B | Carbohydrates | Di- and oligosaccharides | Lactose and Galactose Uptake and Utilization | Tagatose-6-phosphate kinase (EC 2.7.1.144) | no | yes |
| B | Carbohydrates | Di- and oligosaccharides | Sucrose utilization | Sucrose permease, major facilitator superfamily | no | yes |
| B | Carbohydrates | Fermentation | Acetoin, butanediol metabolism | 2,3-butanediol dehydrogenase, S-alcohol forming, (S)-acetoin-specific (EC 1.1.1.76) | no | yes |
| B | Carbohydrates | Monosaccharides | D-Galacturonate and D-Glucuronate Utilization | Pectate lyase precursor (EC 4.2.2.2) | no | yes |
| B | Carbohydrates | Monosaccharides | D-gluconate and ketogluconates metabolism | L-idonate 5-dehydrogenase (EC 1.1.1.264) | no | yes |
| B | Carbohydrates | Monosaccharides | L-Arabinose utilization | Arabinose-proton symporter | no | yes |
| B | Carbohydrates | Monosaccharides | L-Arabinose utilization | L-arabinose isomerase (EC 5.3.1.4) | no | yes |
| B | Carbohydrates | Monosaccharides | L-Arabinose utilization | L-ribulose-5-phosphate 4-epimerase (EC 5.1.3.4) | no | yes |
| B | Carbohydrates | Monosaccharides | L-Arabinose utilization | Ribulokinase (EC 2.7.1.16) | no | yes |
| B | Carbohydrates | Monosaccharides | L-Arabinose utilization | Transcriptional repressor of arabinoside utilization operon, GntR family | no | yes |
| B | Carbohydrates | Sugar alcohols | Inositol catabolism | Inosose isomerase (EC 5.3.99.-) | no | yes |
| B | Cell Wall and Capsule | Capsular and extracellular polysacchrides | Rhamnose containing glycans | Alpha-D-GlcNAc alpha-1,2-L-rhamnosyltransferase (EC 2.4.1.-) | no | yes |
| B | DNA Metabolism | CRISPs | CRISPRs | CRISPR-associated helicase Cas3 | no | yes |
| B | DNA Metabolism | DNA repair | DNA repair, bacterial | ADA regulatory protein | no | yes |
| B | DNA Metabolism | DNA repair | DNA repair, bacterial MutL-MutS system | MutS domain protein, family 4 | no | yes |
| B | DNA Metabolism | no subcategory | Restriction-Modification System | Type III restriction-modification system DNA endonuclease res (EC 3.1.21.5) | no | yes |
| B | DNA Metabolism | no subcategory | Restriction-Modification System | Type III restriction-modification system methylation subunit (EC 2.1.1.72) | no | yes |

**Table S2B.** Functional comparison of differences between the strains TMW 1.1827 vs AGA58 (B=TMW 1.1827, continues)

| **Presence** | **Category** | **Subcategory** | **Subsystem** | **Role** | **SS active A** | **SS active B** |
| --- | --- | --- | --- | --- | --- | --- |
| B | Fatty Acids, Lipids, and Isoprenoids | Fatty acids | Fatty Acid Biosynthesis FASII | Enoyl-[acyl-carrier-protein] reductase [FMN] (EC 1.3.1.9) | no | yes |
| B | Nucleosides and Nucleotides | no subcategory | Hydantoin metabolism | N-carbamoyl-L-amino acid hydrolase (EC 3.5.1.87) | no | yes |
| B | Regulation and Cell signaling | Programmed Cell Death and Toxin-antitoxin Systems | Toxin-antitoxin replicon stabilization systems | YoeB toxin protein | no | yes |
| B | Regulation and Cell signaling | no subcategory | LysR-family proteins in Escherichia coli | Chromosome initiation inhibitor | no | yes |
| B | Stress Response | no subcategory | Carbon Starvation | Starvation sensing protein RspA | no | yes |

**Table S3.** The Protein-BLAST results of the members of the Bacteriocin gene cluster of *Liquorilactobacillus nagelii* AGA58 were predicted using the BAGEL4 webserver.

| **#** | **Name** | **Gene Start (bp)** | **Gene End (bp)** | **Gene strand** | **Description** | **E-Value** | **Percent Identity** | **Accession** |
| --- | --- | --- | --- | --- | --- | --- | --- | --- |
| 1 | orf00001 | 421 | 699 | + | type II toxin-antitoxin system Phd/YefM family antitoxin [*Lactobacillaceae*] | 2e-59 | 100% | WP_027822197.1 |
| 2 | orf00002 | 699 | 1055 | + | type II toxin-antitoxin system YafQ family toxin [*Lactobacillaceae*] | 3e-80 | 100% | WP_027822196.1 |
| 3 | orf00003 | 1258 | 1896 | + | helix-turn-helix domain containing protein [*Liquorilactobacillus hordei*] | 3e-150 | 100% | WP_057870269.1 |
| 4 | Hly D | 2309 | 3694 | - | bacteriocin ABC superfamily ATP binding cassette transporter [*Lactobacillus hordei* DSM 19519] | 0.0 | 99% | KRL05022.1 |
| 5 | Lan T | 3701 | 5830 | - | competence factor transporting permease ATP-binding protein [*Lactobacillus hordei* DSM 19519] | 0.0 | 100% | KRL05021.1 |
| 6 | orf00008 | 6090 | 6299 | + | **Blp family class II bacteriocin [*Liquorilactobacillus hordei*]** | 1e-39 | 100% | WP_057870266.1 |
| 7 | orf00009 | 6314 | 6520 | + | **Blp family class II bacteriocin [*Liquorilactobacillus hordei*]** | 3e-40 | 100% | WP_057870265.1 |
| 8 | orf00010 | 6608 | 7123 | - | DUF3278 domain-containing protein [*Liquorilactobacillus hordei*] | 4e-117 | 100% | WP_057870264.1 |
| 9 | orf00015 | 7658 | 8188 | + | PedC/BrcD family bacteriocin maturation disulfide isomerase [*Liquorilactobacillus hordei*] | 2e-124 | 100% | WP_157047980.1 |
| 10 | EntA_Immun | 8178 | 8558 | + | bacteriocin immunity protein [*Liquorilactobacillus hordei*] | 2e-84 | 100% | WP_157047979.1 |
| 11 | **166.2;Plantaricin_423 (*plaA*)** | 8583 | 8753 | + | **type A2 lantipeptide [*Liquorilactobacillus hordei*]** | 1e-32 | 100% | WP_057870260.1 |
| 12 | orf00019 | 9200 | 9697 | - | helix-turn-helix domain-containing protein [*Liquorilactobacillus hordei*] | 5e-98 | 100% | WP_057870259.1 |
| 13 | orf00021 | 9904 | 10731 | - | hypothetical protein FC81_GL000016 [*Lactobacillus capillatus* DSM 19910] | 0.0 | 99% | KRL03365.1 |
| 14 | orf00022 | 11641 | 12324 | - | IS6 family transposase [*Lactobacillaceae*] | 5e-165 | 100% | WP_022669498.1 |
| 15 | orf00024 | 12321 | 12584 | - | hypothetical protein [*Lactiplantibacillus plantarum*] | 5e-55 | 100% | KZU19941.1 |

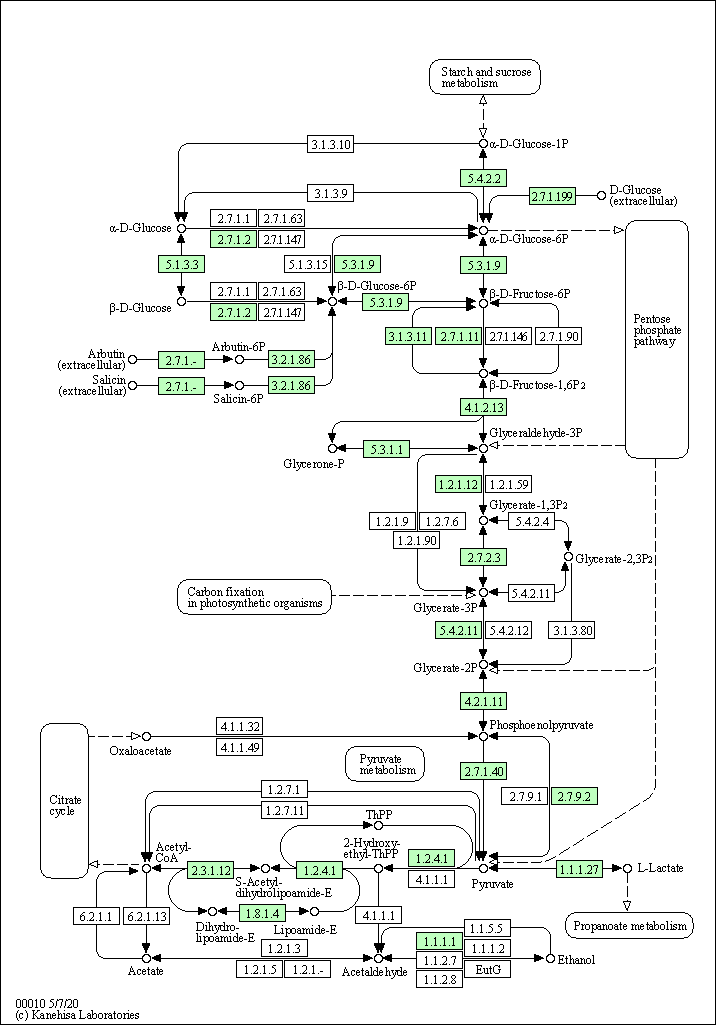

**Fig S1.** The graphical presentation of the possessed enzymes in glycolysis/gluconeogenesis pathways of *Liquorilactobacillus nagelii* AGA58 was obtained from KEGG Mapper (Green coloured EC numbers indicates the presence of the pathway enzymes).

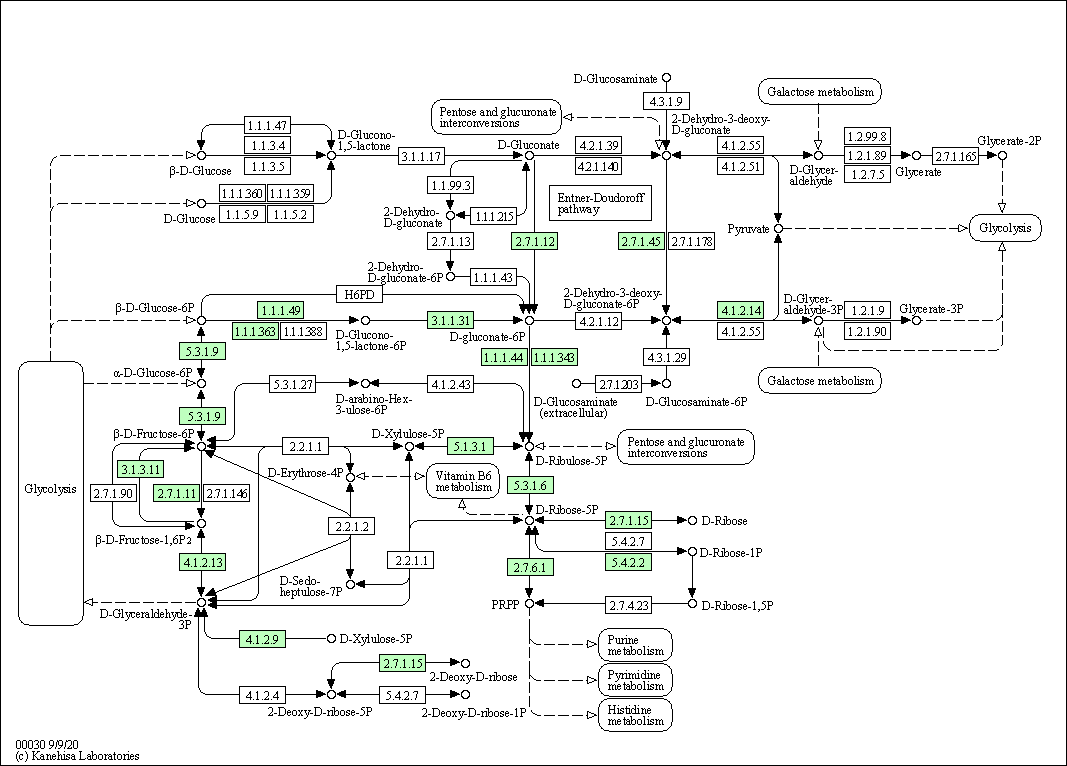

**Fig S2.** The graphical presentation of pentose phosphate pathway enzymes of *Liquorilactobacillus nagelii* AGA 58 was obtained from KEGG Mapper (Green coloured EC number indicates the presence of the pathway enzymes).

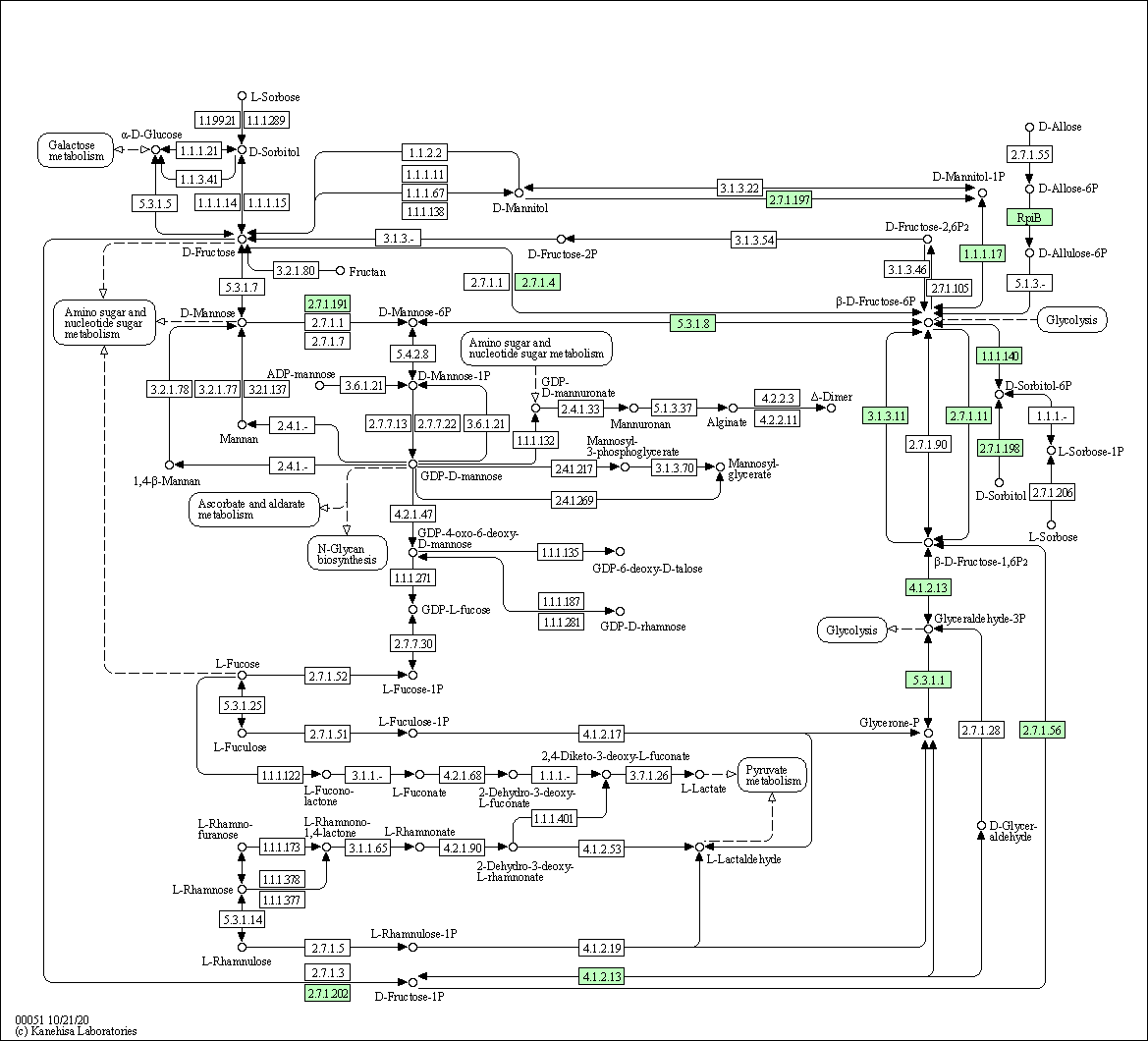

**Fig S3.** The graphical presentation of fructose metabolism pathways of *Liquorilactobacillus nagelii* AGA 58 was obtained from KEGG Mapper (Green coloured EC number indicates the presence of the pathway enzymes).

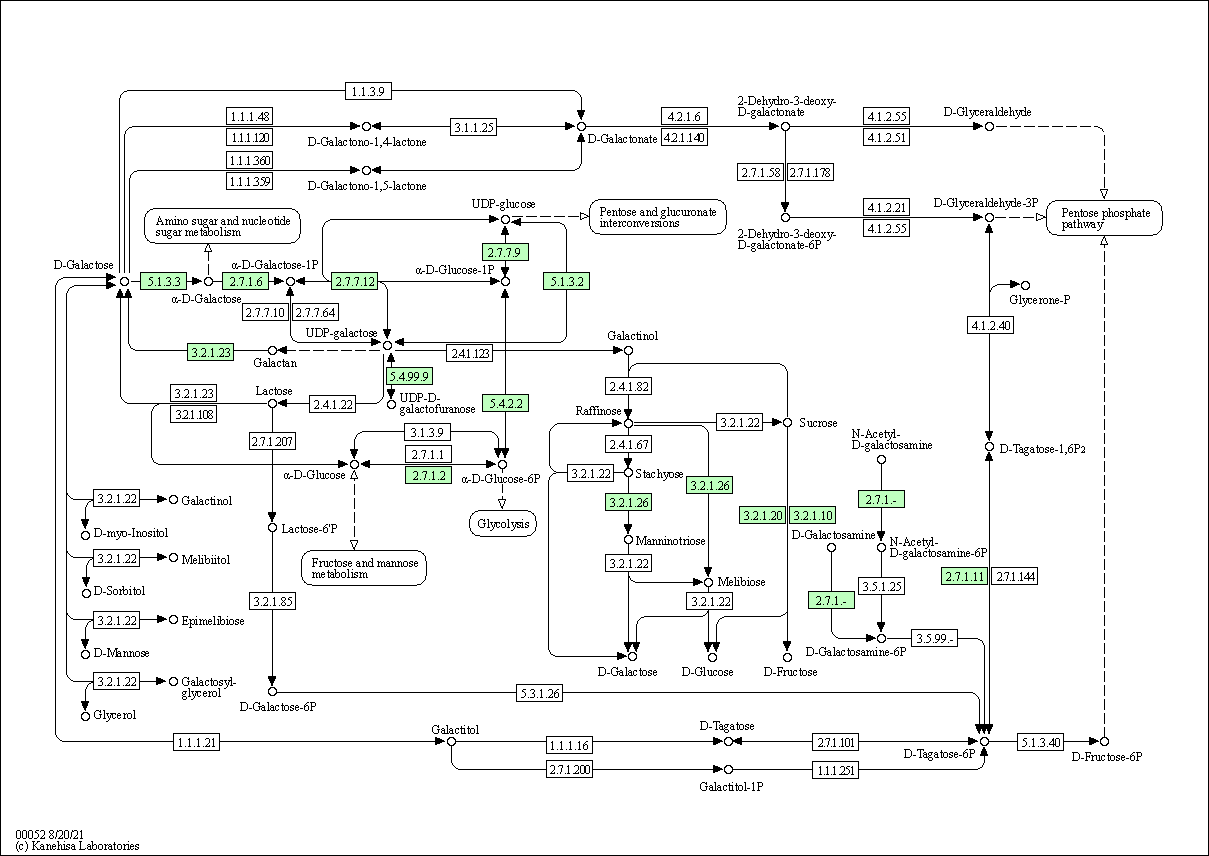

Fig S4. The graphical presentation of galactose metabolism pathways of Liquorilactobacillus nagelii AGA 58 was obtained from KEGG Mapper (Green coloured EC number indicates the presence of the pathway enzyme

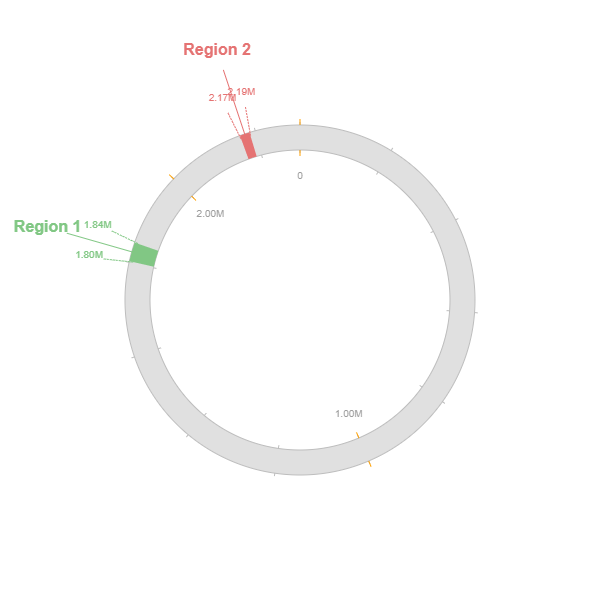
**Fig S3.** The locations of prophage regions on the genome of *Liquorilactobacillus nagelii* AGA58 (green marked region: intact and the score >90, and the red-coloured region: incomplete and the score <70)

**Table S4.** KEGG (BlastKOALA) Orthology search results for the enzymes responsible for carbohydrate metabolism.

| **Glycolysis / Gluconeogenesis** | | | |  |
| --- | --- | --- | --- | --- |
| **#** | **KEGG Entry** | **Symbol** | **Definition** | **Copy Number** |
| 1 | K00016 | LDH, ldh | L-lactate dehydrogenase [EC:1.1.1.27] | 3 |
| 2 | K00134 | GAPDH, gapA | glyceraldehyde 3-phosphate dehydrogenase (phosphorylating) [EC:1.2.1.12] | 1 |
| 3 | K00161 | PDHA, pdhA | pyruvate dehydrogenase E1 component alpha subunit [EC:1.2.4.1] | 1 |
| 4 | K00162 | PDHB, pdhB | pyruvate dehydrogenase E1 component beta subunit [EC:1.2.4.1] | 1 |
| 5 | K00382 | DLD, lpd, pdhD | dihydrolipoamide dehydrogenase [EC:1.8.1.4] | 1 |
| 6 | K00627 | DLAT, aceF, pdhC | pyruvate dehydrogenase E2 component (dihydrolipoamide acetyltransferase) [EC:2.3.1.12] | 1 |
| 7 | K00850 | pfkA, PFK | 6-phosphofructokinase 1 [EC:2.7.1.11] | 1 |
| 8 | K00873 | PK, pyk | pyruvate kinase [EC:2.7.1.40] | 1 |
| 9 | K00927 | PGK, pgk | phosphoglycerate kinase [EC:2.7.2.3] | 1 |
| 10 | K01007 | pps, ppsA | pyruvate, water dikinase [EC:2.7.9.2] | 1 |
| 11 | K01223 | E3.2.1.86B, bglA | 6-phospho-beta-glucosidase [EC:3.2.1.86] | 7 |
| 12 | K01624 | FBA, fbaA | fructose-bisphosphate aldolase, class II [EC:4.1.2.13] | 1 |
| 13 | K01689 | ENO, eno | enolase [EC:4.2.1.11] | 2 |
| 14 | K01785 | galM, GALM | aldose 1-epimerase [EC:5.1.3.3] | 1 |
| 15 | K01803 | TPI, tpiA | triosephosphate isomerase (TIM) [EC:5.3.1.1] | 1 |
| 16 | K01810 | GPI, pgi | glucose-6-phosphate isomerase [EC:5.3.1.9] | 1 |
| 17 | K01834 | PGAM, gpmA | 2,3-bisphosphoglycerate-dependent phosphoglycerate mutase [EC:5.4.2.11] | 2 |
| 18 | K01835 | pgm | phosphoglucomutase [EC:5.4.2.2] | 1 |
| 19 | K02777 | crr | sugar PTS system EIIA component [EC:2.7.1.-] | 1 |
| 20 | K04041 | fbp3 | fructose-1,6-bisphosphatase III [EC:3.1.3.11] | 1 |
| 21 | K04072 | adhE | acetaldehyde dehydrogenase / alcohol dehydrogenase [EC:1.2.1.10 1.1.1.1] | 1 |
| 22 | K25026 | glk | glucokinase [EC:2.7.1.2] | 1 |
| **Citrate cycle (TCA cycle)** | | | |  |
| **#** | **KEGG Entry** | **Symbol** | **Definition** | **Copy Number** |
| 1 | K00031 | IDH1, IDH2, icd | isocitrate dehydrogenase [EC:1.1.1.42] | 1 |
| 2 | K00161 | PDHA, pdhA | pyruvate dehydrogenase E1 component alpha subunit [EC:1.2.4.1] | 1 |
| 3 | K00162 | PDHB, pdhB | pyruvate dehydrogenase E1 component beta subunit [EC:1.2.4.1] | 1 |
| 4 | K00244 | frdA | fumarate reductase flavoprotein subunit [EC:1.3.5.4] | 2 |
| 5 | K00382 | DLD, lpd, pdhD | dihydrolipoamide dehydrogenase [EC:1.8.1.4] | 1 |
| 6 | K00627 | DLAT, aceF, pdhC | pyruvate dehydrogenase E2 component (dihydrolipoamide acetyltransferase) [EC:2.3.1.12] | 1 |
| 7 | K01647 | CS, gltA | citrate synthase [EC:2.3.3.1] | 1 |
| 8 | K01679 | E4.2.1.2B, fumC, FH | fumarate hydratase, class II [EC:4.2.1.2] | 1 |
| 9 | K01681 | ACO, acnA | aconitate hydratase [EC:4.2.1.3] | 1 |
| 10 | K01958 | PC, pyc | pyruvate carboxylase [EC:6.4.1.1] | 1 |

**Table S4.** KEGG (BlastKOALA) Orthology search results for the enzymes responsible for carbohydrate metabolism (continues).

| **Pentose phosphate pathway** | | | |  |
| --- | --- | --- | --- | --- |
| **#** | **KEGG Entry** | **Symbol** | **Definition** | **Copy Number** |
| 1 | K00033 | PGD, gnd, gntZ | 6-phosphogluconate dehydrogenase [EC:1.1.1.44 1.1.1.343] | 2 |
| 2 | K00036 | G6PD, zwf | glucose-6-phosphate 1-dehydrogenase [EC:1.1.1.49 1.1.1.363] | 1 |
| 3 | K00850 | pfkA, PFK | 6-phosphofructokinase 1 [EC:2.7.1.11] | 1 |
| 4 | K00852 | rbsK, RBKS | ribokinase [EC:2.7.1.15] | 1 |
| 5 | K00874 | kdgK | 2-dehydro-3-deoxygluconokinase [EC:2.7.1.45] | 4 |
| 6 | K00948 | PRPS, prsA | ribose-phosphate pyrophosphokinase [EC:2.7.6.1] | 2 |
| 7 | K01621 | xfp, xpk | xylulose-5-phosphate/fructose-6-phosphate phosphoketolase [EC:4.1.2.9 4.1.2.22] | 1 |
| 8 | K01624 | FBA, fbaA | fructose-bisphosphate aldolase, class II [EC:4.1.2.13] | 1 |
| 9 | K01625 | eda | 2-dehydro-3-deoxyphosphogluconate aldolase / (4S)-4-hydroxy-2-oxoglutarate aldolase [EC:4.1.2.14 4.1.3.42] | 2 |
| 10 | K01783 | rpe, RPE | ribulose-phosphate 3-epimerase [EC:5.1.3.1] | 1 |
| 11 | K01807 | rpiA | ribose 5-phosphate isomerase A [EC:5.3.1.6] | 2 |
| 12 | K01808 | rpiB | ribose 5-phosphate isomerase B [EC:5.3.1.6] | 1 |
| 13 | K01810 | GPI, pgi | glucose-6-phosphate isomerase [EC:5.3.1.9] | 1 |
| 14 | K01835 | pgm | phosphoglucomutase [EC:5.4.2.2] | 1 |
| 15 | K04041 | fbp3 | fructose-1,6-bisphosphatase III [EC:3.1.3.11] | 1 |
| 16 | K07404 | pgl | 6-phosphogluconolactonase [EC:3.1.1.31] | 1 |
| 17 | K25031 | gntK | gluconokinase [EC:2.7.1.12] | 1 |
| **Pentose and glucuronate interconversions** | | | |  |
| **#** | **KEGG Entry** | **Symbol** | **Definition** | **Copy Number** |
| 1 | K00040 | uxuB | fructuronate reductase [EC:1.1.1.57] | 1 |
| 2 | K00041 | uxaB | tagaturonate reductase [EC:1.1.1.58] | 1 |
| 3 | K00963 | UGP2, galU, galF | UTP--glucose-1-phosphate uridylyltransferase [EC:2.7.7.9] | 1 |
| 4 | K01685 | uxaA | altronate hydrolase [EC:4.2.1.7] | 1 |
| 5 | K01686 | uxuA | mannonate dehydratase [EC:4.2.1.8] | 1 |
| 6 | K01783 | rpe, RPE | ribulose-phosphate 3-epimerase [EC:5.1.3.1] | 1 |
| 7 | K01812 | uxaC | glucuronate isomerase [EC:5.3.1.12] | 2 |

**Table S4.** KEGG (BlastKOALA) Orthology search results for the enzymes responsible for carbohydrate metabolism (continues).

| **Fructose and mannose metabolism** | | | |  |
| --- | --- | --- | --- | --- |
| **#** | **KEGG Entry** | **Symbol** | **Definition** | **Copy Number** |
| 1 | K00009 | mtlD | mannitol-1-phosphate 5-dehydrogenase [EC:1.1.1.17] | 1 |
| 2 | K00068 | srlD | sorbitol-6-phosphate 2-dehydrogenase [EC:1.1.1.140] | 1 |
| 3 | K00847 | scrK | fructokinase [EC:2.7.1.4] | 1 |
| 4 | K00850 | pfkA, PFK | 6-phosphofructokinase 1 [EC:2.7.1.11] | 1 |
| 5 | K00882 | fruK | 1-phosphofructokinase [EC:2.7.1.56] | 1 |
| 6 | K01624 | FBA, fbaA | fructose-bisphosphate aldolase, class II [EC:4.1.2.13] | 1 |
| 7 | K01803 | TPI, tpiA | triosephosphate isomerase (TIM) [EC:5.3.1.1] | 1 |
| 8 | K01808 | rpiB | ribose 5-phosphate isomerase B [EC:5.3.1.6] | 1 |
| 9 | K01809 | manA, MPI | mannose-6-phosphate isomerase [EC:5.3.1.8] | 1 |
| 10 | K02770 | fruA | fructose PTS system EIIBC or EIIC component [EC:2.7.1.202] | 2 |
| 11 | K02781 | srlB | glucitol/sorbitol PTS system EIIA component [EC:2.7.1.198] | 2 |
| 12 | K02782 | srlE | glucitol/sorbitol PTS system EIIB component [EC:2.7.1.198] | 1 |
| 13 | K02783 | srlA | glucitol/sorbitol PTS system EIIC component | 1 |
| 14 | K02793 | manXa | mannose PTS system EIIA component [EC:2.7.1.191] | 1 |
| 15 | K02794 | manX | mannose PTS system EIIAB component [EC:2.7.1.191] | 2 |
| 16 | K02795 | manY | mannose PTS system EIIC component | 3 |
| 17 | K02796 | manZ | mannose PTS system EIID component | 2 |
| 18 | K02798 | cmtB | mannitol PTS system EIIA component [EC:2.7.1.197] | 1 |
| 19 | K02800 | mtlA, cmtA | mannitol PTS system EIICBA or EIICB component [EC:2.7.1.197] | 1 |
| 20 | K04041 | fbp3 | fructose-1,6-bisphosphatase III [EC:3.1.3.11] | 1 |
| **Galactose metabolism** | | | |  |
| **#** | **KEGG Entry** | **Symbol** | **Definition** | **Copy Number** |
| 1 | K00849 | galK | galactokinase [EC:2.7.1.6] | 1 |
| 2 | K00850 | pfkA, PFK | 6-phosphofructokinase 1 [EC:2.7.1.11] | 1 |
| 3 | K00963 | UGP2, galU, galF | UTP--glucose-1-phosphate uridylyltransferase [EC:2.7.7.9] | 1 |
| 4 | K00965 | galT, GALT | UDPglucose--hexose-1-phosphate uridylyltransferase [EC:2.7.7.12] | 1 |
| 5 | K01182 | IMA, malL | oligo-1,6-glucosidase [EC:3.2.1.10] | 2 |
| 6 | K01187 | malZ | alpha-glucosidase [EC:3.2.1.20] | 1 |
| 7 | K01193 | INV, sacA | beta-fructofuranosidase [EC:3.2.1.26] | 2 |
| 8 | K01784 | galE, GALE | UDP-glucose 4-epimerase [EC:5.1.3.2] | 1 |
| 9 | K01785 | galM, GALM | aldose 1-epimerase [EC:5.1.3.3] | 1 |
| 10 | K01835 | pgm | phosphoglucomutase [EC:5.4.2.2] | 1 |
| 11 | K01854 | glf | UDP-galactopyranose mutase [EC:5.4.99.9] | 3 |
| 12 | K02744 | agaF | N-acetylgalactosamine PTS system EIIA component [EC:2.7.1.-] | 1 |
| 13 | K12308 | bgaB, lacA | beta-galactosidase [EC:3.2.1.23] | 1 |
| 14 | K25026 | glk | glucokinase [EC:2.7.1.2] | 1 |

**Table S4.** KEGG (BlastKOALA) Orthology search results for the enzymes responsible for carbohydrate metabolism (continues).

| **Starch and sucrose metabolism** | | | |  |
| --- | --- | --- | --- | --- |
| **#** | **KEGG Entry** | **Symbol** | **Definition** | **Copy Number** |
| 1 | K00688 | PYG, glgP | glycogen phosphorylase [EC:2.4.1.1] | 1 |
| 2 | K00700 | GBE1, glgB | 1,4-alpha-glucan branching enzyme [EC:2.4.1.18] | 1 |
| 3 | K00703 | glgA | starch synthase [EC:2.4.1.21] | 1 |
| 4 | K00847 | E2.7.1.4, scrK | fructokinase [EC:2.7.1.4] | 1 |
| 5 | K00963 | UGP2, galU, galF | UTP--glucose-1-phosphate uridylyltransferase [EC:2.7.7.9] | 1 |
| 6 | K00975 | glgC | glucose-1-phosphate adenylyltransferase [EC:2.7.7.27] | 2 |
| 7 | K01182 | IMA, malL | oligo-1,6-glucosidase [EC:3.2.1.10] | 2 |
| 8 | K01187 | malZ | alpha-glucosidase [EC:3.2.1.20] | 1 |
| 9 | K01193 | INV, sacA | beta-fructofuranosidase [EC:3.2.1.26] | 2 |
| 10 | K01208 | cd, ma, nplT | cyclomaltodextrinase / maltogenic alpha-amylase / neopullulanase [EC:3.2.1.54 3.2.1.133 3.2.1.135] | 1 |
| 11 | K01223 | E3.2.1.86B, bglA | 6-phospho-beta-glucosidase [EC:3.2.1.86] | 7 |
| 12 | K01226 | treC | trehalose-6-phosphate hydrolase [EC:3.2.1.93] | 1 |
| 13 | K01810 | GPI, pgi | glucose-6-phosphate isomerase [EC:5.3.1.9] | 1 |
| 14 | K01835 | pgm | phosphoglucomutase [EC:5.4.2.2] | 1 |
| 15 | K02759 | celC, chbA | cellobiose PTS system EIIA component [EC:2.7.1.196 2.7.1.205] | 2 |
| 16 | K02760 | celA, chbB | cellobiose PTS system EIIB component [EC:2.7.1.196 2.7.1.205] | 2 |
| 17 | K02761 | celB, chbC | cellobiose PTS system EIIC component | 3 |
| 18 | K02777 | crr | sugar PTS system EIIA component [EC:2.7.1.-] | 1 |
| 19 | K02810 | scrA, sacP, sacX, ptsS | sucrose PTS system EIIBCA or EIIBC component [EC:2.7.1.211] | 1 |
| 20 | K25026 | glk | glucokinase [EC:2.7.1.2] | 1 |

**Table S4.** KEGG (BlastKOALA) Orthology search results for the enzymes responsible for carbohydrate metabolism (continues).

| **Amino sugar and nucleotide sugar metabolism** | | | |  |
| --- | --- | --- | --- | --- |
| **#** | **KEGG Entry** | **Symbol** | **Definition** | **Copy Number** |
| 1 | K00075 | murB | UDP-N-acetylmuramate dehydrogenase [EC:1.3.1.98] | 1 |
| 2 | K00790 | murA | UDP-N-acetylglucosamine 1-carboxyvinyltransferase [EC:2.5.1.7] | 2 |
| 3 | K00820 | glmS, GFPT | glutamine---fructose-6-phosphate transaminase (isomerizing) [EC:2.6.1.16] | 1 |
| 4 | K00847 | E2.7.1.4, scrK | fructokinase [EC:2.7.1.4] | 1 |
| 5 | K00849 | galK | galactokinase [EC:2.7.1.6] | 1 |
| 6 | K00963 | UGP2, galU, galF | UTP--glucose-1-phosphate uridylyltransferase [EC:2.7.7.9] | 1 |
| 7 | K00965 | galT, GALT | UDPglucose--hexose-1-phosphate uridylyltransferase [EC:2.7.7.12] | 1 |
| 8 | K00975 | glgC | glucose-1-phosphate adenylyltransferase [EC:2.7.7.27] | 2 |
| 9 | K01207 | nagZ | beta-N-acetylhexosaminidase [EC:3.2.1.52] | 1 |
| 10 | K01443 | nagA, AMDHD2 | N-acetylglucosamine-6-phosphate deacetylase [EC:3.5.1.25] | 1 |
| 11 | K01784 | galE, GALE | UDP-glucose 4-epimerase [EC:5.1.3.2] | 1 |
| 12 | K01791 | wecB | UDP-N-acetylglucosamine 2-epimerase (non-hydrolysing) [EC:5.1.3.14] | 1 |
| 13 | K01809 | manA, MPI | mannose-6-phosphate isomerase [EC:5.3.1.8] | 1 |
| 14 | K01810 | GPI, pgi | glucose-6-phosphate isomerase [EC:5.3.1.9] | 1 |
| 15 | K01835 | pgm | phosphoglucomutase [EC:5.4.2.2] | 1 |
| 16 | K01854 | glf | UDP-galactopyranose mutase [EC:5.4.99.9] | 3 |
| 17 | K02564 | nagB, GNPDA | glucosamine-6-phosphate deaminase [EC:3.5.99.6] | 1 |
| 18 | K02777 | crr | sugar PTS system EIIA component [EC:2.7.1.-] | 1 |
| 19 | K02793 | manXa | mannose PTS system EIIA component [EC:2.7.1.191] | 1 |
| 20 | K02794 | manX | mannose PTS system EIIAB component [EC:2.7.1.191] | 2 |
| 21 | K02795 | manY | mannose PTS system EIIC component | 3 |
| 22 | K02796 | manZ | mannose PTS system EIID component | 2 |
| 23 | K03431 | glmM | phosphoglucosamine mutase [EC:5.4.2.10] | 1 |
| 24 | K04042 | glmU | bifunctional UDP-N-acetylglucosamine pyrophosphorylase / glucosamine-1-phosphate N-acetyltransferase [EC:2.7.7.23 2.3.1.157] | 1 |
| 25 | K07106 | murQ | N-acetylmuramic acid 6-phosphate etherase [EC:4.2.1.126] | 1 |
| 26 | K25026 | glk | glucokinase [EC:2.7.1.2] | 1 |

**Table S4.** KEGG (BlastKOALA) Orthology search results for the enzymes responsible for carbohydrate metabolism (continues).

| **Pyruvate metabolism** | | | |  |
| --- | --- | --- | --- | --- |
| **#** | **KEGG Entry** | **Symbol** | **Definition** | **Copy Number** |
| 1 | K00016 | LDH, ldh | L-lactate dehydrogenase [EC:1.1.1.27] | 3 |
| 2 | K00027 | ME2, sfcA, maeA | malate dehydrogenase (oxaloacetate-decarboxylating) [EC:1.1.1.38] | 2 |
| 3 | K00158 | E1.2.3.3, poxL | pyruvate oxidase [EC:1.2.3.3] | 2 |
| 4 | K00161 | PDHA, pdhA | pyruvate dehydrogenase E1 component alpha subunit [EC:1.2.4.1] | 1 |
| 5 | K00162 | PDHB, pdhB | pyruvate dehydrogenase E1 component beta subunit [EC:1.2.4.1] | 1 |
| 6 | K00244 | frdA | fumarate reductase flavoprotein subunit [EC:1.3.5.4] | 2 |
| 7 | K00382 | DLD, lpd, pdhD | dihydrolipoamide dehydrogenase [EC:1.8.1.4] | 1 |
| 8 | K00625 | E2.3.1.8, pta | phosphate acetyltransferase [EC:2.3.1.8] | 1 |
| 9 | K00626 | ACAT, atoB | acetyl-CoA C-acetyltransferase [EC:2.3.1.9] | 1 |
| 10 | K00627 | DLAT, aceF, pdhC | pyruvate dehydrogenase E2 component (dihydrolipoamide acetyltransferase) [EC:2.3.1.12] | 1 |
| 11 | K00656 | E2.3.1.54, pflD | formate C-acetyltransferase [EC:2.3.1.54] | 1 |
| 12 | K00873 | PK, pyk | pyruvate kinase [EC:2.7.1.40] | 1 |
| 13 | K00925 | ackA | acetate kinase [EC:2.7.2.1] | 4 |
| 14 | K01007 | pps, ppsA | pyruvate, water dikinase [EC:2.7.9.2] | 1 |
| 15 | K01512 | acyP | acylphosphatase [EC:3.6.1.7] | 1 |
| 16 | K01649 | leuA, IMS | 2-isopropylmalate synthase [EC:2.3.3.13] | 1 |
| 17 | K01679 | E4.2.1.2B, fumC, FH | fumarate hydratase, class II [EC:4.2.1.2] | 1 |
| 18 | K01958 | PC, pyc | pyruvate carboxylase [EC:6.4.1.1] | 1 |
| 19 | K01961 | accC | acetyl-CoA carboxylase, biotin carboxylase subunit [EC:6.4.1.2 6.3.4.14] | 2 |
| 20 | K01962 | accA | acetyl-CoA carboxylase carboxyl transferase subunit alpha [EC:6.4.1.2 2.1.3.15] | 1 |
| 21 | K01963 | accD | acetyl-CoA carboxylase carboxyl transferase subunit beta [EC:6.4.1.2 2.1.3.15] | 1 |
| 22 | K02160 | accB, bccP | acetyl-CoA carboxylase biotin carboxyl carrier protein | 2 |
| 23 | K03778 | ldhA | D-lactate dehydrogenase [EC:1.1.1.28] | 2 |
| 24 | K04072 | adhE | acetaldehyde dehydrogenase / alcohol dehydrogenase [EC:1.2.1.10 1.1.1.1] | 1 |
| 25 | K22212 | mleA, mleS | malolactic enzyme [EC:4.1.1.101] | 2 |
| 26 | K22373 | larA | lactate racemase [EC:5.1.2.1] | 2 |

**Table S4** KEGG (BlastKOALA) Orthology search results for the enzymes responsible for carbohydrate metabolism (continues).

| **Glyoxylate and dicarboxylate metabolism** | | | |  |
| --- | --- | --- | --- | --- |
| **#** | **KEGG Entry** | **Symbol** | **Definition** | **Copy Number** |
| 1 | K00018 | hprA | glycerate dehydrogenase [EC:1.1.1.29] | 1 |
| 2 | K00382 | DLD, lpd, pdhD | dihydrolipoamide dehydrogenase [EC:1.8.1.4] | 1 |
| 3 | K00600 | glyA, SHMT | glycine hydroxymethyltransferase [EC:2.1.2.1] | 1 |
| 4 | K00626 | ACAT, atoB | acetyl-CoA C-acetyltransferase [EC:2.3.1.9] | 1 |
| 5 | K00865 | glxK, garK | glycerate 2-kinase [EC:2.7.1.165] | 1 |
| 6 | K01091 | gph | phosphoglycolate phosphatase [EC:3.1.3.18] | 1 |
| 7 | K01625 | eda | 2-dehydro-3-deoxyphosphogluconate aldolase / (4S)-4-hydroxy-2-oxoglutarate aldolase [EC:4.1.2.14 4.1.3.42] | 2 |
| 8 | K01647 | CS, gltA | citrate synthase [EC:2.3.3.1] | 1 |
| 9 | K01681 | ACO, acnA | aconitate hydratase [EC:4.2.1.3] | 1 |
| 10 | K01915 | glnA, GLUL | glutamine synthetase [EC:6.3.1.2] | 1 |
| **Propanoate metabolism** | | | |  |
| **#** | **KEGG Entry** | **Symbol** | **Definition** | **Copy Number** |
| 1 | K00016 | LDH, ldh | L-lactate dehydrogenase [EC:1.1.1.27] | 3 |
| 2 | K00086 | dhaT | 1,3-propanediol dehydrogenase [EC:1.1.1.202] | 1 |
| 3 | K00382 | DLD, lpd, pdhD | dihydrolipoamide dehydrogenase [EC:1.8.1.4] | 1 |
| 4 | K00625 | E2.3.1.8, pta | phosphate acetyltransferase [EC:2.3.1.8] | 1 |
| 5 | K00656 | E2.3.1.54, pflD | formate C-acetyltransferase [EC:2.3.1.54] | 1 |
| 6 | K00925 | ackA | acetate kinase [EC:2.7.2.1] | 4 |
| 7 | K01961 | accC | acetyl-CoA carboxylase, biotin carboxylase subunit [EC:6.4.1.2 6.3.4.14] | 2 |
| 8 | K01962 | accA | acetyl-CoA carboxylase carboxyl transferase subunit alpha [EC:6.4.1.2 2.1.3.15] | 1 |
| 9 | K01963 | accD | acetyl-CoA carboxylase carboxyl transferase subunit beta [EC:6.4.1.2 2.1.3.15] | 1 |
| 10 | K02160 | accB, bccP | acetyl-CoA carboxylase biotin carboxyl carrier protein | 2 |

**Table S4** KEGG (BlastKOALA) Orthology search results for the enzymes responsible for carbohydrate metabolism (continues).

| **Butanoate metabolism** | | | |  |
| --- | --- | --- | --- | --- |
| **#** | **KEGG Entry** | **Symbol** | **Definition** | **Copy Number** |
| 1 | K00004 | BDH, butB | (R,R)-butanediol dehydrogenase / meso-butanediol dehydrogenase / diacetyl reductase [EC:1.1.1.4 1.1.1.- 1.1.1.303] | 2 |
| 2 | K00022 | HADH | 3-hydroxyacyl-CoA dehydrogenase [EC:1.1.1.35] | 1 |
| 3 | K00135 | gabD | succinate-semialdehyde dehydrogenase / glutarate-semialdehyde dehydrogenase [EC:1.2.1.16 1.2.1.79 1.2.1.20] | 2 |
| 4 | K00244 | frdA | fumarate reductase flavoprotein subunit [EC:1.3.5.4] | 2 |
| 5 | K00626 | ACAT, atoB | acetyl-CoA C-acetyltransferase [EC:2.3.1.9] | 1 |
| 6 | K00656 | E2.3.1.54, pflD | formate C-acetyltransferase [EC:2.3.1.54] | 1 |
| 7 | K01575 | alsD, budA, aldC | acetolactate decarboxylase [EC:4.1.1.5] | 1 |
| 8 | K01641 | HMGCS | hydroxymethylglutaryl-CoA synthase [EC:2.3.3.10] | 1 |
| 9 | K01652 | E2.2.1.6L, ilvB, ilvG, ilvI | acetolactate synthase I/II/III large subunit [EC:2.2.1.6] | 2 |
| 10 | K04072 | adhE | acetaldehyde dehydrogenase / alcohol dehydrogenase [EC:1.2.1.10 1.1.1.1] | 1 |
| 11 | K18120 | 4hbD, abfH | 4-hydroxybutyrate dehydrogenase [EC:1.1.1.61] | 1 |
| **C5-Branched dibasic acid metabolism** | | | |  |
| **#** | **KEGG Entry** | **Symbol** | **Definition** | **Copy Number** |
| 1 | K00052 | leuB, IMDH | 3-isopropylmalate dehydrogenase [EC:1.1.1.85] | 1 |
| 2 | K01575 | alsD, budA, aldC | acetolactate decarboxylase [EC:4.1.1.5] | 1 |
| 3 | K01652 | E2.2.1.6L, ilvB, ilvG, ilvI | acetolactate synthase I/II/III large subunit [EC:2.2.1.6] | 2 |
| 4 | K01703 | leuC, IPMI-L | 3-isopropylmalate/(R)-2-methylmalate dehydratase large subunit [EC:4.2.1.33 4.2.1.35] | 1 |
| 5 | K01704 | leuD, IPMI-S | 3-isopropylmalate/(R)-2-methylmalate dehydratase small subunit [EC:4.2.1.33 4.2.1.35] | 1 |
| **Inositol phosphate metabolism** | | | |  |
| **#** | **KEGG Entry** | **Symbol** | **Definition** | **Copy Number** |
| 1 | K00010 | iolG | myo-inositol 2-dehydrogenase / D-chiro-inositol 1-dehydrogenase [EC:1.1.1.18 1.1.1.369] | 3 |
| 2 | K01092 | E3.1.3.25, IMPA, suhB | myo-inositol-1(or 4)-monophosphatase [EC:3.1.3.25] | 1 |
| 3 | K01803 | TPI, tpiA | triosephosphate isomerase (TIM) [EC:5.3.1.1] | 1 |

**Table S5.** KEGG (BlastKOALA) Orthology search results for ABC transporters and phosphotransferase system (PTS).

| **ABC Transporters** | | | |  |
| --- | --- | --- | --- | --- |
| **#** | **KEGG Entry** | **Symbol** | **Definition** | **Copy Number** |
| 1 | K02036 | pstB | phosphate transport system ATP-binding protein [EC:7.3.2.1] | 3 |
| 2 | K02037 | pstC | phosphate transport system permease protein | 2 |
| 3 | K02038 | pstA | phosphate transport system permease protein | 2 |
| 4 | K02040 | pstS | phosphate transport system substrate-binding protein | 2 |
| 5 | K02071 | metN | D-methionine transport system ATP-binding protein | 1 |
| 6 | K02072 | metI | D-methionine transport system permease protein | 1 |
| 7 | K02073 | metQ | D-methionine transport system substrate-binding protein | 1 |
| 8 | K02424 | fliY, tcyA | L-cystine transport system substrate-binding protein | 1 |
| 9 | K03523 | bioY | biotin transport system substrate-specific component | 1 |
| 10 | K05813 | ugpB | sn-glycerol 3-phosphate transport system substrate-binding protein | 1 |
| 11 | K05814 | ugpA | sn-glycerol 3-phosphate transport system permease protein | 1 |
| 12 | K05815 | ugpE | sn-glycerol 3-phosphate transport system permease protein | 1 |
| 13 | K05816 | ugpC | sn-glycerol 3-phosphate transport system ATP-binding protein [EC:7.6.2.10] | 1 |
| 14 | K05845 | opuC | osmoprotectant transport system substrate-binding protein | 1 |
| 15 | K05846 | opuBD | osmoprotectant transport system permease protein | 2 |
| 16 | K05847 | opuA | osmoprotectant transport system ATP-binding protein [EC:7.6.2.9] | 1 |
| 17 | K09693 | tagH | teichoic acid transport system ATP-binding protein [EC:7.5.2.4] | 1 |
| 18 | K09811 | ftsX | cell division transport system permease protein | 1 |
| 19 | K09812 | ftsE | cell division transport system ATP-binding protein | 2 |
| 20 | K10009 | tcyB, yecS | L-cystine transport system permease protein | 1 |
| 21 | K10010 | tcyC, yecC | L-cystine transport system ATP-binding protein [EC:7.4.2.1] | 1 |
| 22 | K10036 | glnH | glutamine transport system substrate-binding protein | 2 |
| 23 | K10037 | glnP | glutamine transport system permease protein | 2 |
| 24 | K10038 | glnQ | glutamine transport system ATP-binding protein [EC:7.4.2.1] | 1 |
| 25 | K10823 | oppF | oligopeptide transport system ATP-binding protein | 1 |
| 26 | K15580 | oppA, mppA | oligopeptide transport system substrate-binding protein | 7 |
| 27 | K15581 | oppB | oligopeptide transport system permease protein | 1 |
| 28 | K15582 | oppC | oligopeptide transport system permease protein | 1 |

**Table S5.** KEGG (BlastKOALA) Orthology search results for ABC transporters and phosphotransferase system (PTS) (Continues).

| **ABC Transporters** | | | |  |
| --- | --- | --- | --- | --- |
| **#** | **KEGG Entry** | **Symbol** | **Definition** | **Copy Number** |
| 29 | K15583 | oppD | oligopeptide transport system ATP-binding protein | 1 |
| 30 | K16012 | cydC | ATP-binding cassette, subfamily C, bacterial CydC | 1 |
| 31 | K16013 | cydD | ATP-binding cassette, subfamily C, bacterial CydD | 1 |
| 32 | K16785 | ecfT | energy-coupling factor transport system permease protein | 3 |
| 33 | K16786 | ecfA1 | energy-coupling factor transport system ATP-binding protein [EC:7.-.-.-] | 1 |
| 34 | K16787 | ecfA2 | energy-coupling factor transport system ATP-binding protein [EC:7.-.-.-] | 1 |
| 35 | K17077 | artQ | arginine/lysine/histidine transport system permease protein | 1 |
| 36 | K18887 | efrA, efrE | ATP-binding cassette, subfamily B, multidrug efflux pump | 1 |
| 37 | K18888 | efrB, efrF | ATP-binding cassette, subfamily B, multidrug efflux pump | 1 |
| 38 | K18891 | patA, rscA, lmrC, satA | ATP-binding cassette, subfamily B, multidrug efflux pump | 1 |
| 39 | K18892 | patB, rscB, lmrC, satB | ATP-binding cassette, subfamily B, multidrug efflux pump | 1 |
| 40 | K19083 | braD, bceA | bacitracin transport system ATP-binding protein | 1 |
| 41 | K19084 | braE, bceB | bacitracin transport system permease protein | 1 |
| 42 | K20344 | blpA, lagD | ATP-binding cassette, subfamily C, bacteriocin exporter | 1 |
| 43 | K23059 | artP, artI | arginine/lysine/histidine transporter system substrate-binding protein | 1 |
| 44 | K23060 | artR, artM | arginine/lysine/histidine transport system ATP-binding protein [EC:7.4.2.1] | 1 |

**Table S5.** KEGG (BlastKOALA) Orthology search results for ABC transporters and phosphotransferase system (PTS) (Continues).

| **Phosphotransferase system (PTS)** | | | |  |
| --- | --- | --- | --- | --- |
| **#** | **KEGG Entry** | **Symbol** | **Definition** | **Copy Number** |
| 1 | K00882 | fruK | 1-phosphofructokinase [EC:2.7.1.56] | 1 |
| 2 | K02744 | agaF | N-acetylgalactosamine PTS system EIIA component [EC:2.7.1.-] | 1 |
| 3 | K02757 | bglF, bglP | beta-glucoside PTS system EIICBA component [EC:2.7.1.-] | 3 |
| 4 | K02759 | celC, chbA | cellobiose PTS system EIIA component [EC:2.7.1.196 2.7.1.205] | 2 |
| 5 | K02760 | celA, chbB | cellobiose PTS system EIIB component [EC:2.7.1.196 2.7.1.205] | 2 |
| 6 | K02761 | celB, chbC | cellobiose PTS system EIIC component | 3 |
| 7 | K02770 | fruA | fructose PTS system EIIBC or EIIC component [EC:2.7.1.202] | 2 |
| 8 | K02777 | crr | sugar PTS system EIIA component [EC:2.7.1.-] | 1 |
| 9 | K02781 | srlB | glucitol/sorbitol PTS system EIIA component [EC:2.7.1.198] | 2 |
| 10 | K02782 | srlE | glucitol/sorbitol PTS system EIIB component [EC:2.7.1.198] | 1 |
| 11 | K02783 | srlA | glucitol/sorbitol PTS system EIIC component | 1 |
| 12 | K02784 | ptsH | phosphocarrier protein HPr | 1 |
| 13 | K02793 | manXa | mannose PTS system EIIA component [EC:2.7.1.191] | 1 |
| 14 | K02794 | manX | mannose PTS system EIIAB component [EC:2.7.1.191] | 2 |
| 15 | K02795 | manY | mannose PTS system EIIC component | 3 |
| 16 | K02796 | manZ | mannose PTS system EIID component | 2 |
| 17 | K02798 | cmtB | mannitol PTS system EIIA component [EC:2.7.1.197] | 1 |
| 18 | K02800 | mtlA, cmtA | mannitol PTS system EIICBA or EIICB component [EC:2.7.1.197] | 1 |
| 19 | K02810 | scrA, sacP, sacX, ptsS | sucrose PTS system EIIBCA or EIIBC component [EC:2.7.1.211] | 1 |
| 20 | K08483 | ptsI | phosphoenolpyruvate-protein phosphotransferase (PTS system enzyme I) [EC:2.7.3.9] | 1 |

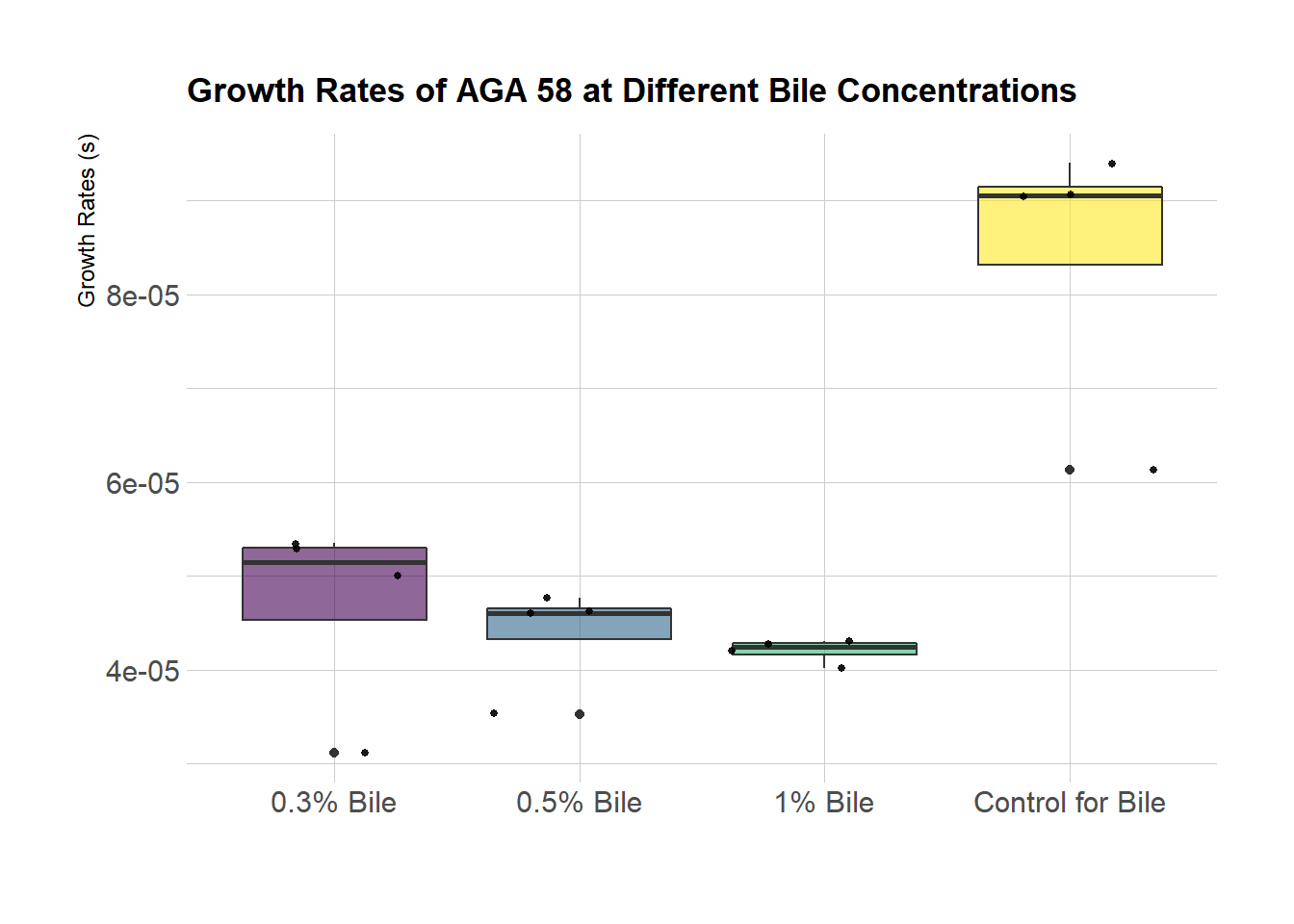

**Fig S4.** Calculated growth rates when *L. nagelii* was grown in control, 0.3%, 0.5% or 1% bile salt conditions

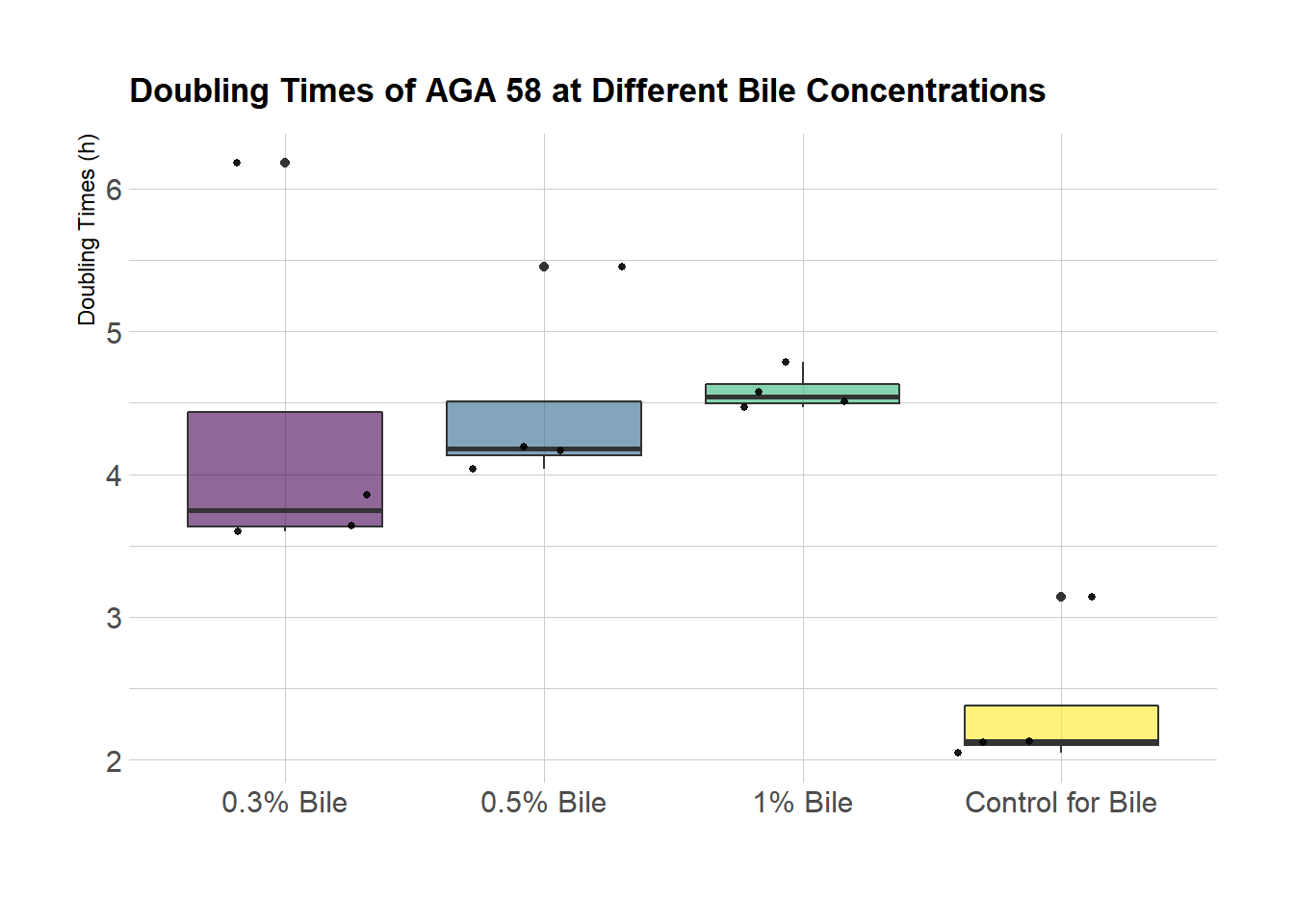

**Fig S5.** Calculated doubling times calculated when *L. nagelii* was grown in control, 0.3%, 0.5% or 1% bile salt conditions

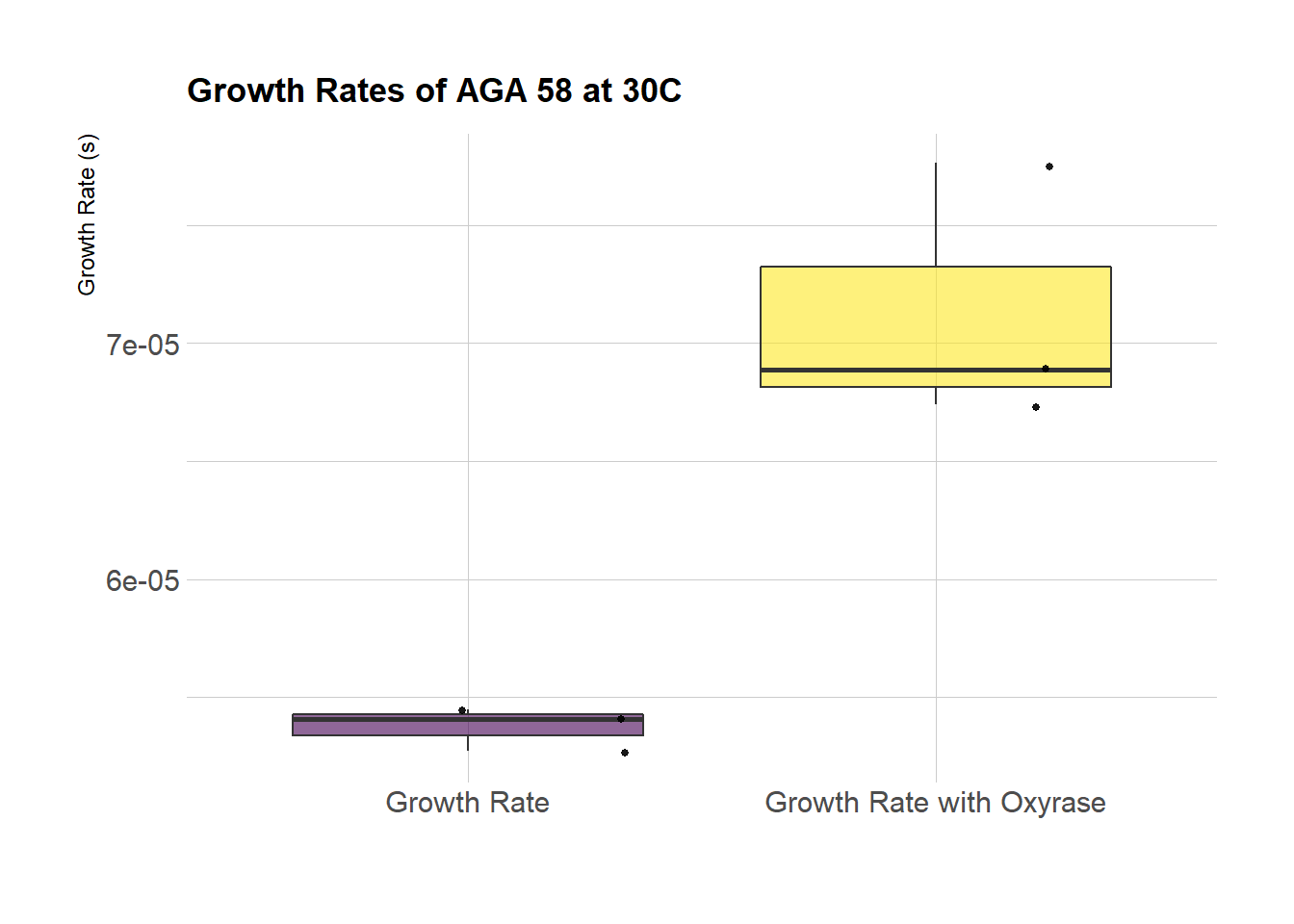

**Fig S6.** Calculated growth rates when *L. nagelii* AGA58 was grown with added oxyrase

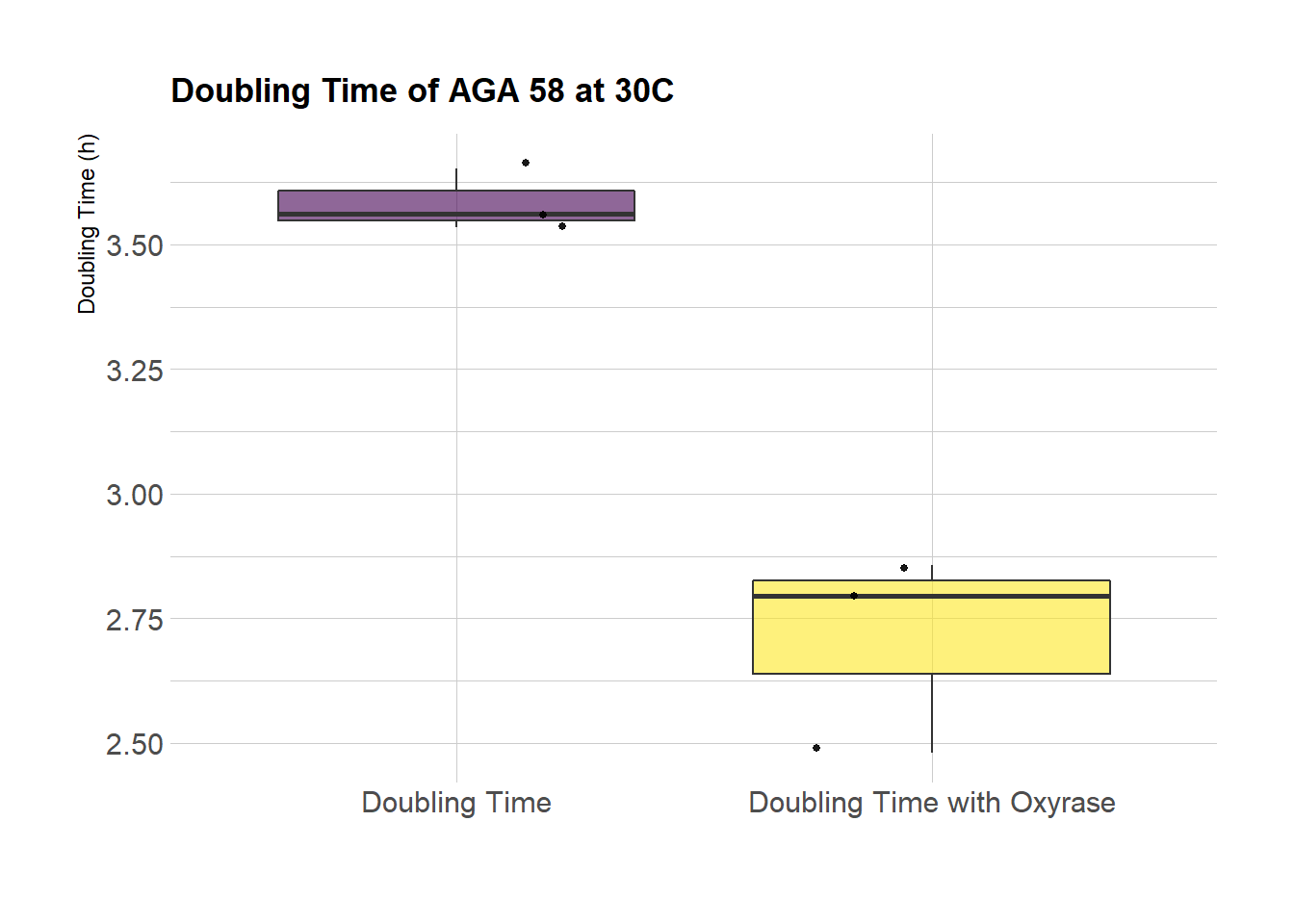

**Fig S7.** Calculated doubling times when *L. nagelii* AGA58 was grown with added oxyrase

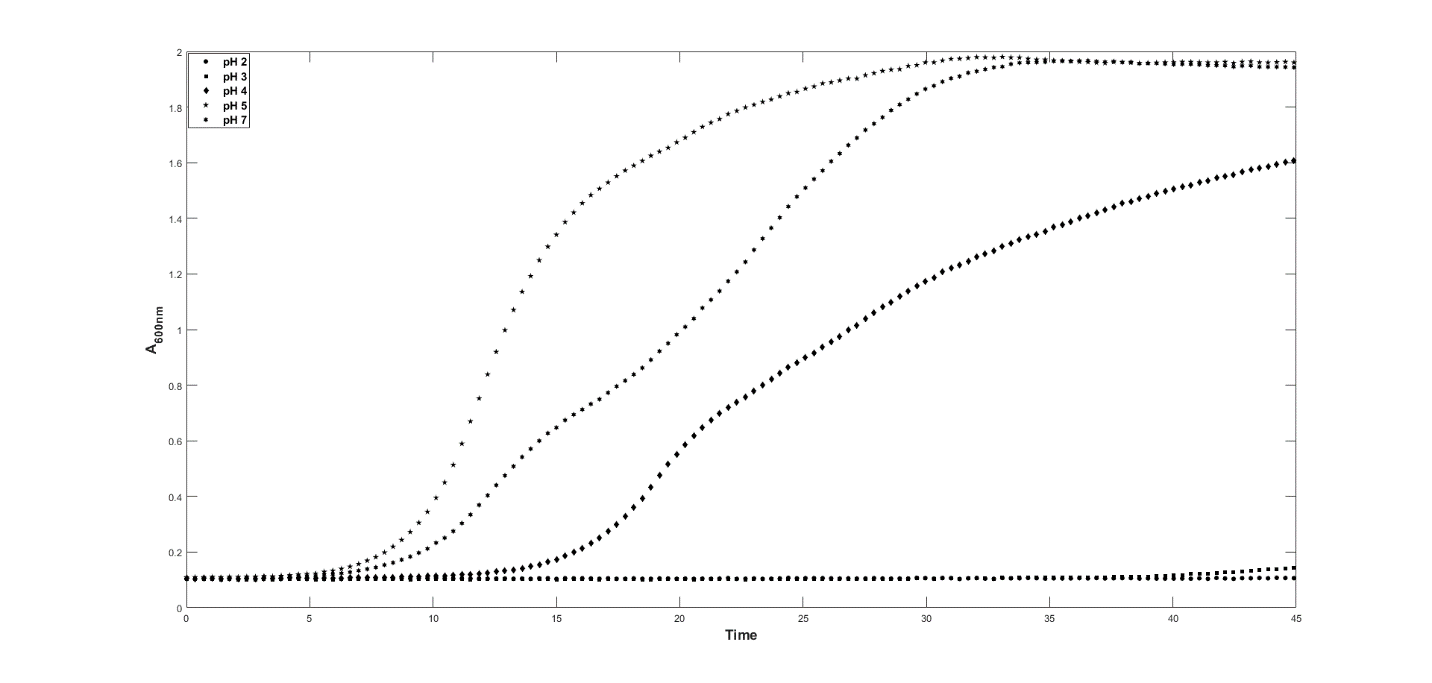

**Fig S6.** The growth curves at pH 2, 3, 5 and 7 for *L. nagelii* AGA 58

**Table S7.** KEGG orthology (KO) functional categories of identified protein-coding sequences in the genome of *Liquorilactobacillus nagelii* AGA58

| **KO Number** | **Functional category** | **Gene Number** | **Proportion (%)** |
| --- | --- | --- | --- |
| 09101 | Carbohydrate metabolism | 173 | 12,53 |
| 09102 | Energy metabolism | 29 | 2,10 |
| 09103 | Lipid metabolism | 35 | 2,53 |
| 09104 | Nucleotide metabolism | 58 | 4,20 |
| 09105 | Amino acid metabolism | 90 | 6,52 |
| 09106 | Metabolism of other amino acids | 14 | 1,01 |
| 09107 | Glycan biosynthesis and metabolism | 22 | 1,59 |
| 09108 | Metabolism of cofactors and vitamins | 48 | 3,48 |
| 09110 | Metabolism of terpenoids and polyketides | 8 | 0,58 |
| 09111 | Xenobiotics biodegradation and metabolism | 3 | 0,22 |
| 09120 | Genetic information processing | 153 | 11,08 |
| 09130 | Environmental information processing | 111 | 8,04 |
| 09140 | Cellular processes | 46 | 3,33 |
| 09150 | Organismal systems | 6 | 0,43 |
| 09181 | Protein families: metabolism | 39 | 2,82 |
| 09182 | Protein families: genetic information processing | 188 | 13,61 |
| 09183 | Protein families: signalling and cellular processes | 144 | 10,43 |
| 09191 | Unclassified: metabolism | 84 | 6,08 |
| 09192 | Unclassified: genetic information processing | 50 | 3,62 |
| 09193 | Unclassified: signaling and cellular processes | 25 | 1,81 |
| - | Unclassified | 54 | 3,91 |
